# SMN promotes mitochondrial metabolic maturation during myogenesis by regulating the MYOD-miRNA axis

**DOI:** 10.1101/2022.02.13.480288

**Authors:** Akihiro Ikenaka, Yohko Kitagawa, Michiko Yoshida, Chuang-Yu Lin, Akira Niwa, Tatsutoshi Nakahata, Megumu K. Saito

## Abstract

Spinal muscular atrophy (SMA) is a congenital neuromuscular disease caused by the mutation or deletion of *survival motor neuron 1* (*SMN1*) gene. Although the primary cause of progressive muscle atrophy in SMA has classically been considered the degeneration of motor neurons, recent studies have indicated a skeletal muscle-specific pathological phenotype such as impaired mitochondrial function and enhanced cell death. Here we found that the downregulation of SMN causes mitochondrial dysfunction and subsequent cell death in *in vitro* models of skeletal myogenesis with both a murine C2C12 cell line and human induced pluripotent stem cells. During myogenesis, SMN binds to the genome upstream of the transcriptional start sites of MYOD1 and microRNA (miR)-1 and -206. Accordingly, the loss of SMN downregulates these miRs, whereas supplementation of the miRs recovers the mitochondrial function, cell survival and myotube formation of SMN-deficient C2C12, indicating the SMN-miR axis is essential for myogenic metabolic maturation. Additionally, introduction of the miRs into ex vivo muscle stem cells derived from Δ7-SMA mice caused myotube formation and muscle contraction. In conclusion, our data revealed novel transcriptional roles of SMN during myogenesis, providing an alternative muscle-oriented therapeutic strategy for SMA patients.

**Highlights:**

- Reduced SMN causes mitochondrial dysregulation in myogenic cells.
- Reduced SMN downregulates miR-1 and miR-206 expression in myogenic cells.
- SMN protein binds to the genome upstream of MYOD1, miR-1 and miR-206.
- miR-1 and miR-206 are sufficient to improve skeletal muscle function in an SMA model.

## Introduction

Spinal muscular atrophy (SMA) is an inherent neuromuscular disease caused by mutation or deletion of *survival motor neuron 1* (*SMN1*) gene. *SMN1* encodes the SMN protein. In the severest form of SMA, infants suffer from severe muscle weakness and respiratory failure during the first year of life. Classically, reduced SMN expression was thought to cause selective and primary lower motor neuronal death, leading to subsequent denervation and muscle atrophy, since the degeneration of anterior horn motor neurons is the predominant pathological finding of SMA. However, SMA is currently regarded as a systemic disorder affecting not only motor neurons, but also neuromuscular junctions and skeletal muscles (Hayhurst et al., 2012; Martinez et al., 2012). Indeed, myoblasts with SMN knockdown show reduced proliferation and fusion defects (Shafey et al., 2005). Animal models of SMA also revealed skeletal muscle has its own pathological contribution to the SMA phenotype (Kim et al., 2020; Martinez et al., 2012). However, the cell-autonomous molecular mechanism of the skeletal muscle degeneration due to SMN deficiency is mostly unknown.

SMN localizes ubiquitously and exerts its function in various manners. The first identified function of SMN was the assembly of spliceosomal small nuclear ribonucleoproteins (snRNPs). SMN forms a large protein complex (SMN complex) and chaperones the biogenesis of snRNPs in the cytoplasm and subsequent translocation to the nucleus (Qing Liu, 1997). In neurons, SMN has a unique role on mRNA transport and local translation, which is exerted by binding with HuD in the cytoplasm (Akten et al., 2011; Hao le et al., 2017). SMN is also involved in actin dynamics and axon elongation by interacting with profilin 2a (Akten et al., 2011; Hao le et al., 2017; Sharma et al., 2005). Additionally, the loss of SMN in motor neurons can cause mitochondrial dysfunction and impaired cellular respiration (Miller et al., 2016). These findings indicate that SMN has both universal and cell-type specific roles. Thus, understanding the specific function of SMN in skeletal muscle could lead to novel muscle-targeting therapeutic strategies.

Skeletal muscle stem cells and myoblasts undergo a dramatic bioenergetic transition from glycolysis to oxidative phosphorylation during differentiation into myotubes. This metabolic maturation is accompanied by mitochondrial maturation (Miller et al., 2016; Remels et al., 2010). One of the most important transcription factors governing the metabolic and mitochondrial maturation is *Myoblast determination protein 1* (*MYOD1*). MYOD1 is known to regulate oxidative metabolism by directly binding the enhancers along oxidative metabolic genes (Shintaku et al., 2016). MYOD1 also contributes to metabolic maturation by upregulating microRNA (miR)-1, −133a and −206, all of which are important for metabolic and mitochondrial maturation (Przanowska et al., 2020; Shintaku et al., 2016). The MYOD1-miR axis is therefore important for skeletal muscle biogenesis. Interestingly, skeletal muscle specimens obtained from SMA patients showed a downregulation of electron transfer chain (ETC) and mitochondrial outer membrane proteins (Ripolone et al., 2015), a phenomenon also seen in neurons. Oxygen consumption was also decreased in an *in vitro* model using myotubes differentiated from SMA pluripotent stem cell (PSC) models (Hellbach et al., 2018; Ripolone et al., 2015). Although these findings highlight mitochondrial dysfunction in skeletal muscles, the precise molecular mechanism remains unknown. Especially, the relationship between SMN and the MYOD1-miR axis should be elucidated.

Here we describe a novel function of SMN during myogenesis using human and murine in vitro models. We found that SMN diffusely localizes in the nucleus during myogenesis. The loss of SMN expression causes the downregulation of MYOD1, miR-1 and miR-206, resulting in impaired metabolic maturation and myotube formation. We also show that the *in vitro* phenotypes of SMA myotubes are successfully rescued by the ectopic expression of miRs, demonstrating SMN acts as an upstream regulator of these genes. Interestingly, SMN binds to the upstream genomic legions of MYOD1 and the miR host genes, indicating that SMN regulates their expression. Our results highlight the unique stage- and cell type-specific functions of nuclear SMN that make it a regulatory factor of the metabolic maturation of myotubes, thus providing novel insights into the skeletal myogenesis and an alternative therapeutic strategy for the biogenesis of skeletal muscle in SMA.

## Results

### Loss of SMN causes mitochondrial bioenergetic failure during the myogenic conversion of iPSCs

Knowing that the forced expression of MYOD1 converts human PSCs to myotubes (Tanaka et al., 2013), we first investigated the effect of SMN downregulation on the myogenic conversion of human iPSCs. For this purpose, we prepared two pairs of isogenic iPSC clones with a doxycycline-inducible MYOD1 expression construct: first, a control 201B7 iPSC line and its SMN-knockdown counterpart (B7-M and B7-M^SMNKD^, respectively); and second, an SMA patient-derived iPSC line (Yoshida et al., 2015) and the same line but with SMN supplementation (SMA-M and SMA-M^OE^, respectively). We then converted these clones into myogenic cells by adding doxycycline (**Figure 1A**). On day 3, more than 90% of the cells were positive for Myogenin (MyoG) in all clones (**Figures S1A and S1B**). Genes associated with myogenesis, such as MyoG and Myocyte Enhancer factor 2C (MEF2C), were also upregulated after differentiation (**Figure S1C**). The level of SMN protein was lower in B7-M^SMNKD^ and SMA-M even after differentiation (**Figure S1D**). Interestingly, we found that while iPSC clones with sufficient SMN expression (B7-M and SMA-M^OE^; SMN-maintained clones) showed increased cell number during the conversion, those with downregulated SMN (B7-M^SMNKD^ and SMA-M; SMN-downregulated clones) failed to proliferate (**Figure S1E**). This effect could be attributed to the increased apoptosis in the SMN-downregulated clones, because the cleavage of caspase-3 was increased. Indeed, treatment with a pan-caspase inhibitor, Z-VAD-FMK, reduced the number of cleaved caspase-3 positive cells in SMN-downregulated clones (**Figure 1B and 1C**).

**Figure 1.**
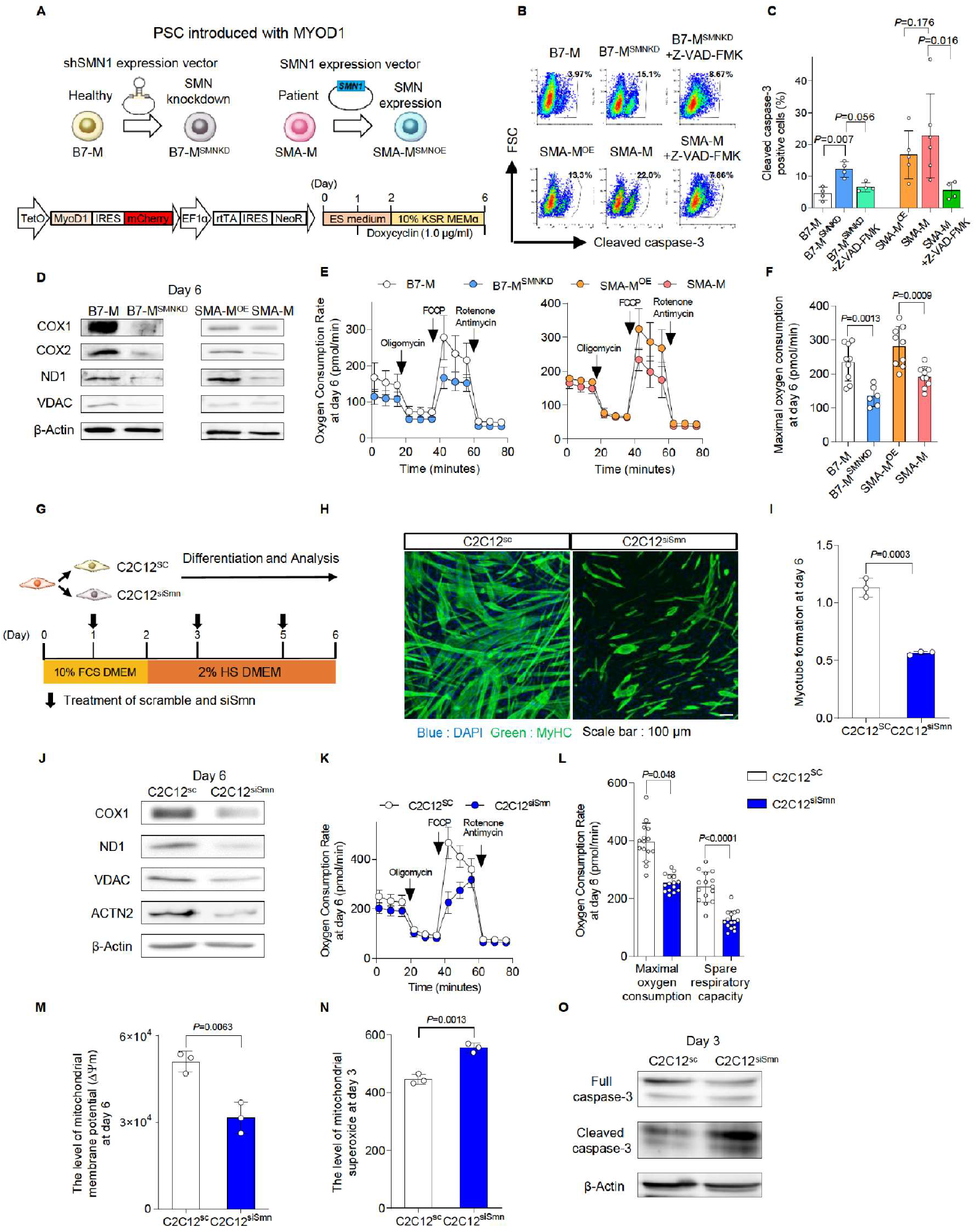
Loss of SMN causes mitochondrial bioenergetic failure during myogenic differentiation. **(A)** The schema of iPSC clones used in the study and the doxycycline-inducible MYOD1-driven myogenic conversion system. **(B, C)** Cleaved caspase-3-positive apoptotic cells in human iPSC-derived myogenic cells (day 3) analyzed by intracellular flow cytometry. Z-VAD-FMK (20 μM) was added. (B) Representative flow diagrams and (C) their quantification. **(D)** Immunoblotting assay with iPSC-derived myogenic cells (day 6). **(E)** OCR measured in iPSC-derived myogenic cells (day 6). Oligomycin (10 μM), FCCP (10 μM) and Rotenone (1.0 μM) plus Antimycin (1.0 μM) were sequentially added. **(F)** Maximal oxygen consumption calculated using the data in (E). **(G)** The schema of myogenic differentiation with C2C12 cells. **(H)** Representative immunostaining images of C2C12 cells with MyHC (myosin heavy chain) (day 6). **(I)** Myotube formation defined by the ratio of the DAPI-positive area to MyHC-positive area (day 6). **(J)** Immunoblotting assay with C2C12 cells (day 6). **(K)** OCR measured with C2C12 cells (day 6). Oligomycin (10 μM), FCCP (10 μM) and Rotenone (1.0 μM) plus Antimycin (1.0 μM) were sequentially added. **(L)** Maximal OCR and spare respiratory capacity calculated using the data in (K). Data are value obtained from 10^4^ cells. **(M)** Mitochondrial membrane potential (Δψm) of C2C12 cells (day 6). **(N)** Mitochondrial superoxide levels of C2C12 cells (day 3) evaluated with MitoSOX. **(O)** Immunoblotting assay with C2C12 cells (day 3). Error bars indicate means ± S.D. (C) Statistical analysis by one-way ANOVA with multiple comparisons. (F, I and K-M) Statistical analysis by Student’s t-test. Each dot represents a biologically independent sample.

During myogenic differentiation, the bioenergetic status shifts from glycolysis to mitochondrial oxidative phosphorylation. To enhance the mitochondrial function, genes associated with the mitochondrial respiratory complex are upregulated (Remels et al., 2010; Wust et al., 2018). To understand the mechanism of apoptosis, we focused on the mitochondrial biology during myogenic conversion, because myogenic differentiation promotes mitochondrial biogenesis (Frangini et al., 2013) and because mitochondrial failure can cause apoptosis. During the myogenic conversion, the mitochondrial DNA (mtDNA) copy number increased in SMN-maintained clones but not in SMN-downregulated clones (**Figure S1F**). SMN-downregulated myogenic cells showed a downregulation of mitochondrial proteins associated with bioenergetic function (**Figure 1D**). The downregulation of mitochondrial proteins generally represses the oxygen consumption rate of mitochondria (Trotta et al., 2017; Yan et al., 2019). Consistently, the mitochondrial oxygen consumption capacity was impaired (**Figures 1E, 1F and S1G**). In line with this finding, the mitochondrial membrane potential (Δψm) was lower in SMN-downregulated clones (**Figure S1H**). These results indicate that SMN is required for mitochondrial bioenergetic maturation during myogenic conversion.

Impairment of the mitochondrial complex in the ETC promotes excessive ROS production, resulting in apoptosis (Trotta et al., 2017). Consistently, ROS production was increased in SMN-downregulated clones on day 3 (**Figure S1I**). Treating the SMN-downregulated clones with an antioxidant, α-tocopherol (αToc), reduced the apoptotic cell number such that it almost equaled that of SMN-maintained clones (**Figures S1J**) and recovered the total cell number (**Figure S1K**). Overall, our observations suggest that the loss of SMN causes mitochondrial bioenergetic dysregulation and subsequent ROS-mediated apoptosis during myogenic conversion.

### Depletion of SMN causes mitochondrial dysfunction in C2C12 cells

Since the iPSC model uses an artificial expression of MYOD1 to convert cell fate, we next evaluated the reproducibility of our findings by employing a commonly used myogenic differentiation model with the murine C2C12 cell line. For this, we knocked down *Smn* in C2C12 (C2C12^siSmn^) and differentiated the cells into myotubes by changing the culture medium (**Figure 1G**). C2C12^siSmn^ showed reduced SMN expression at both the transcript and protein levels (**Figures S1L and S1M**). Differentiated C2C12^siSmn^ showed decreased myotube formation, as measured by myosin heavy chain (MyHC) staining (**Figures 1H and 1I**), indicating the indispensable role of Smn on myogenic differentiation.

The mitochondrial function of C2C12 cells was also evaluated. Cytochrome c oxidase subunit 1 (COX1) was upregulated in C2C12 cells 6 days after the differentiation, indicating mitochondrial maturation (**Figure S1N and S1O)**. On the other hand, mitochondrial proteins in differentiated C2C12^siSmn^ were less compared to C2C12^SC^ (**Figure 1J**). As expected, C2C12^siSmn^ showed a lower oxygen consumption capacity than C2C12^SC^ (**Figures 1K and 1L**), indicating the dysregulation of mitochondrial electron transmission. Consistently, Δψm was also decreased in C2C12^siSmn^ (**Figure 1M**).

As expected, mitochondrial ROS production increased in C2C12^siSmn^ on day 3 of the differentiation (**Figure 1N**). The level of cleaved caspase-3 also increased on day 3 in C2C12^siSmm^, indicating the occurrence of mitochondrial apoptosis (**Figure 1O**). Overall, the data obtained with the C2C12 model were consistent with the iPSC model, showing reproducibility among different species and differentiation systems. Previous SMA studies about the pathogenesis of skeletal muscle showed a failure of the myogenic terminal differentiation, including the abnormal expression of myogenic regulatory genes and fewer myotubes (Boyer et al., 2013; Bricceno et al., 2014; Hayhurst et al., 2012; Nicole et al., 2003). Considering that the failure of the mitochondrial metabolic transition impairs myogenic terminal differentiation (Ripolone et al., 2015; Wust et al., 2018), our results indicate the potential relationship between the mitochondrial metabolic transition and failure of myogenic differentiation in SMA skeletal muscle models.

### Depletion of SMN downregulates the expression of MYOD1 and downstream miR-1 and miR-206

MYOD1 is an essential transcriptional factor for myogenesis, and it and its downstream factors enhance mitochondrial oxidative metabolism during myogenic differentiation (Shintaku et al., 2016; Wust et al., 2018; Zhang et al., 2014). Since the myogenic differentiation of iPSCs, primary myoblasts and C2C12 cells relies on the expression of MYOD1 (Tanaka et al., 2013; Wang et al., 2017), we hypothesized that SMN regulates muscle differentiation via MYOD1. In C2C12 cells, MyoD1 expression increased during differentiation (**Figure S2A**), but the depletion of Smn significantly decreased the expression of Myod1 at both the mRNA and protein level (**Figures 2A and 2B**). Interestingly, in the iPSC model, while the expression of exogenous MYOD1 transgene was comparable between SMN-maintained clones and SMN-downregulated clones (**Figure S2B**), the endogenous MYOD1 expression was significantly impaired in the SMN-downregulated clones (**Figure S2C**). Therefore, the endogenous transcriptional control of MYOD1 gene was affected by the dose of SMN.

**Figure 2.**
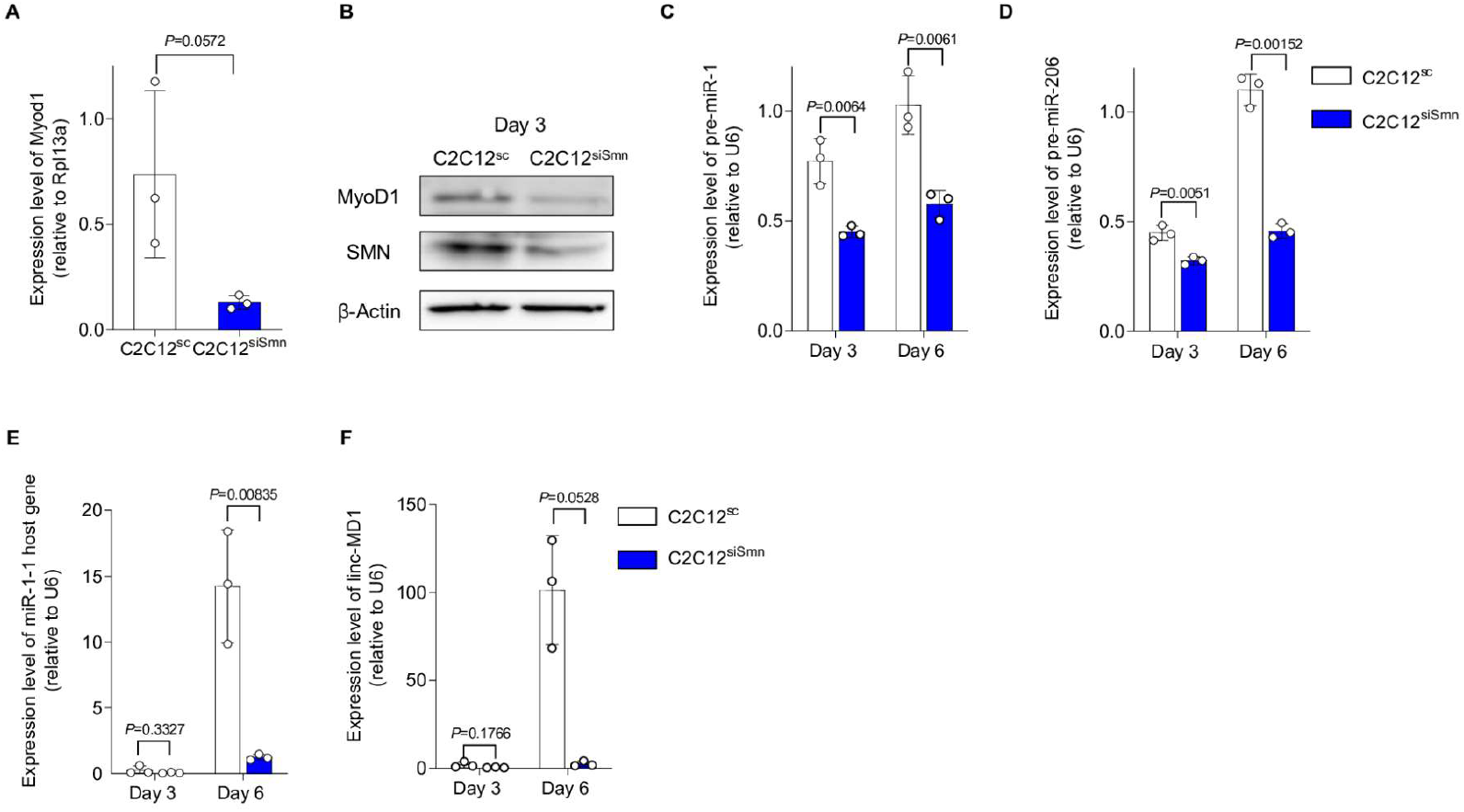
Depletion of SMN downregulates the expression of MYOD1 and its downstream targets miR-1 and miR-206. **(A, B)** Expression of MyoD1 in C2C12^SC^ and C2C12^siSmn^ (day 6) at the (A) mRNA and (B) protein level. (A) Rpl13a served as the internal control. (B) β-Actin was used as the loading control. **(C-F)** qPCR analysis for the expression of (C) pre-miR-1, (D) pre-miR-206, (E) miR1-1 host gene and (F) linc-MD1. U6 served as the internal control. Error bars indicate means ± SD. Statistical analysis by Student’s t-test. Each dot represents a biologically independent sample.

We next investigated a downstream molecular mechanism harnessing MYOD1 and mitochondrial biogenesis. We focused on miR-1 and miR-206, because both are directly regulated by MYOD1, highly expressed during the development of skeletal muscle (Rao et al., 2006), and control mitochondrial function in myogenic cells (Przanowska et al., 2020; Wust et al., 2018; Zhang et al., 2014). As expected, C2C12^siSmn^ showed less expression of miR-1 and miR-206 compared to C2C12^SC^ (**Figures 2C and 2D**). Similarly, in the iPSC model, SMN-downregulated clones showed less expression too (**Figures S2D and S2E**).

Murine miR-1 and miR-206 are embedded in miR1-1 host gene (miR1-1 hg) and long intergenic non-protein coding RNA muscle differentiation 1 (linc-MD1), respectively. These miRs are transcribed as primary miRs (pri-miRs) from these host genes. To determine if the downregulation of the miRs occurred at the transcriptional level, we evaluated the expression of their pri-miRs. The expressions of precursor miRs (pre-miRs) and the host genes of both miRs were also significantly downregulated (**Figures 2E and 2F**), indicating that the cause of impaired miR expression is at least partially attributed to transcriptional dysregulation. In conclusion, our data showed that the depletion of SMN reduces the expression of MYOD1 and its downstream miRs, which in turn could cause impaired mitochondrial metabolic maturation during muscle differentiation.

### Supplementation of miRs to SMN-depleted myoblasts recovers mitochondrial metabolism and myogenesis

To investigate whether miR-1 and miR-206 are responsible factors for the mitochondrial metabolic dysfunction in SMN-depleted myogenic cells, we introduced miR-1 and/or miR-206 into C2C12^siSmn^ before myogenic differentiation (**Figure 3A**). Supplementation of the miRs into C2C12^siSmn^ improved the oxygen consumption capacity on day 6 compared to untreated C2C12^siSmn^ (**Figure 3B and 3C**). Consistently, miR treatment recovered the expression of COX1 and ND1 to the level of C2C12^SC^ (**Figures 3D, 3E and 3F**). The effect of miR supplementation on mitochondrial metabolism was specific to C2C12^siSmn^, as miR treatment to C2C12^SC^ did not affect the mitochondrial oxygen consumption capacity or the expression of COX1 (**Figures S3A, S3B and S3C**). Notably, miR supplementation improved the myogenesis of C2C12^siSmn^, as it recovered the myotube formation ability and cell number (**Figures 3G, 3H and 3I**). In conclusion, the dysregulation of mitochondrial metabolism and myogenesis of myogenic cells due to SMN depletion is caused by the downregulation of miRs.

**Figure 3.**
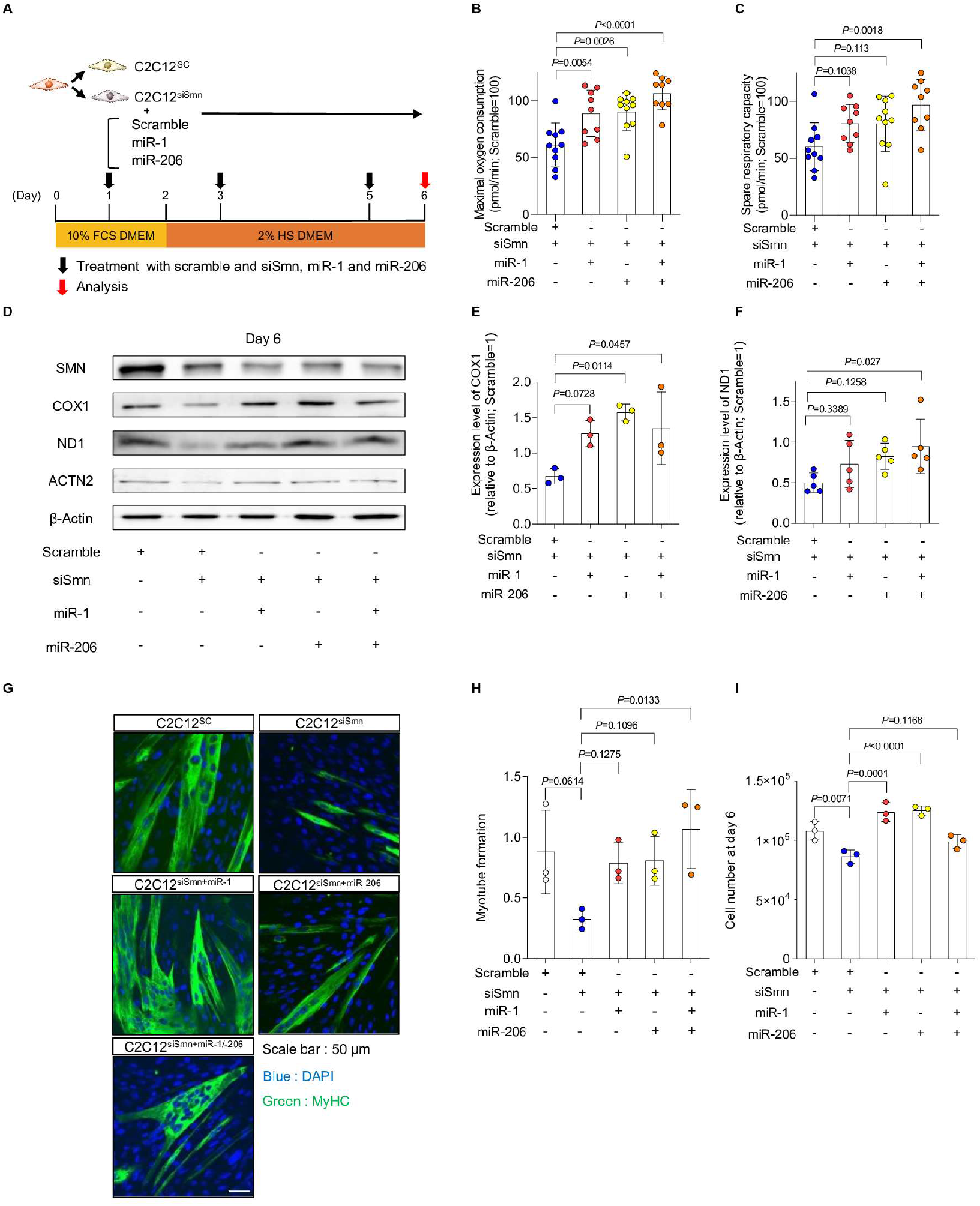
Supplementation of miRNA recovers the phenotypes of SMN-depleted C2C12 cells. **(A)** Schema of the culture system. **(B, C)** (B) Maximal oxygen consumption and (C) spare respiratory capacity of C2C12^siSmn^ cells (day 6). **(D-F)** (D) Immunoblotting assay with C2C12 cells (day 6), and (E, F) quantification of COX1 and ND1 protein. β-Actin served as the loading control. Relative values to C2C12^SC^ (day 6) are shown. **(G)** Representative immunostaining images of C2C12 cells (day 6). **(H)** Myotube formation defined by the ratio of the DAPI-positive area to MyHC-positive area (day 6). **(I)** Number of C2C12 cells (day 6). Error bars indicate means ± SD. Statistical analysis by one-way ANOVA with multiple comparisons. Each dot represents a biologically independent sample.

Next, to address the causal association of the expression of MYOD1 and miRs, exogeneous MYOD1 was introduced into C2C12 cells (**Figure S3D**). MYOD1 overexpression in C2C12^siSmn^ restored both the pri-miR and pre-miR expression to levels comparable with C2C12^SC^ (**Figures S3E and S3F**). It also recovered the expression of proteins associated with mitochondrial metabolism (**Figure S3G**) and myotube formation (**Figures S3H and S3I**). Hence, MYOD1 depletion is responsible for the decrease of these miRs. Overall, our data demonstrate that SMN contributes to the maturation of mitochondrial metabolism and myogenesis by regulating the MYOD1-miR-1/-206 axis.

### SMN transiently localizes in the nucleus during myogenesis

We next sought to understand the molecular mechanisms by which SMN regulates the MYOD1-miR-1/-206 axis. For this purpose, we first tracked the expression and localization of SMN. In the iPSC model, SMN was upregulated at both the transcript and protein level (**Figures 4A, 4B and 4C**), peaking at day 3 of the conversion. Interestingly, we found that SMN protein diffusely localized in the whole nucleus, corresponding to the upregulation of SMN (**Figures 4D and 4E**). The temporal localization of SMN in the nucleus was confirmed by protein extraction of the nuclear compartment (**Figure 4F**). Similar to the iPSC model, the upregulation and diffuse nuclear localization of SMN was observed during the myogenic differentiation of C2C12 cells, which is different from the typical localization of SMN on Cajal bodies (**Figures S4A, S4B, S4C and S4D**). Therefore, we hypothesized that during myogenesis SMN has a unique role in the nucleus.

**Figure 4.**
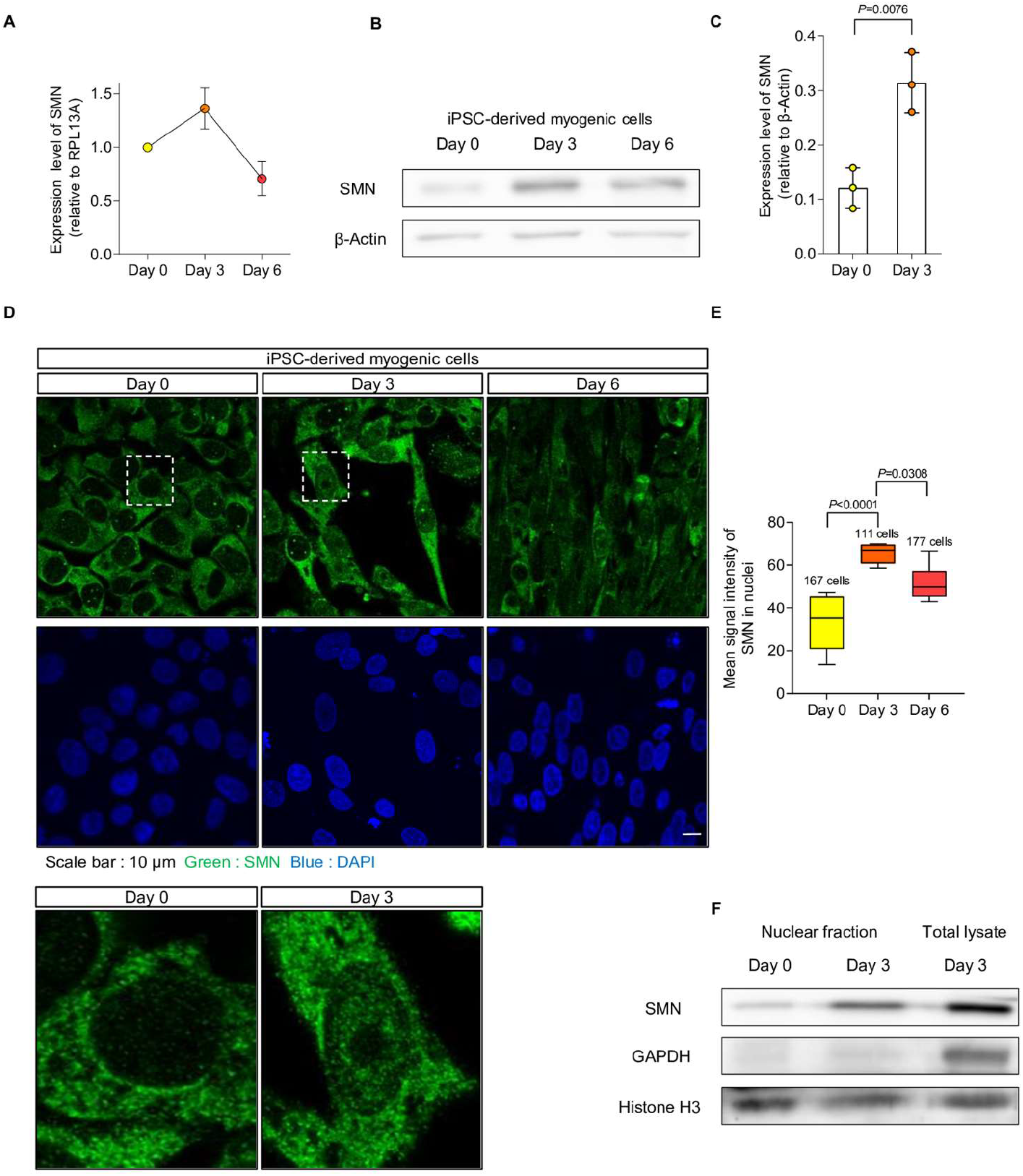
SMN transiently localizes in the nucleus during myogenesis. **(A)** SMN expression in B7-M-derived myogenic cells. Relative values to the expression level at day 0 are plotted. RPL13A was used as the internal control. **(B)** Immunoblotting assay with B7-M-derived myogenic cells. β-Actin served as the loading control. **(C)** Quantification of the SMN expression measured from (B). **(D)** Representative immunostaining images of 201B7 iPSC- and B7-M-derived myogenic cells. Magnified images of the dotted square regions are shown below. **(E)** Quantification of the signal intensity of nuclear SMN using the data in (D). **(F)** Immunoblotting assay with the nuclear fraction separated from 201B7 iPSC- and B7-M-derived myogenic cells. GAPDH and Histone H3 served as the loading control for total lysate and nuclear fraction, respectively. Error bars indicate means ± SD. (C) Statistical analysis by Student’s t-test. (E) Statistical analysis by one-way ANOVA with multiple comparisons. Each dot represents a biologically independent sample.

The canonical function of SMN is to form snRNPs in the cytoplasm and nuclear Cajal bodies in order to maintain the spliceosome (Lotti et al., 2012; Zhang et al., 2008). The number of nuclear SMN foci, which represents the localization of SMN in Cajal bodies, transiently increased in iPSC-derived myogenic cells (**Figure S4E**). However, the signal intensity of diffusely localized SMN protein was not significantly different between cells with and without foci (**Figure S4F**). Furthermore, SMN foci in C2C12 cells did not increase after differentiation despite the nuclear translocation of SMN (**Figure S4G**). These findings imply that diffusely distributed nuclear SMN is regulated independently of SMN in foci and that this spatiotemporally specific distribution is associated with the function of SMN during myogenesis.

### SMN and MYOD1 interact with each other and bind to the promoter regions of myogenic genes

Previous studies reported that SMN is involved in genome instability and transcriptional termination by binding with RNAPL II (RNA polymerase II) (Jangi et al., 2017; Zhao et al., 2016). However, it is unclear whether SMN interacts with other molecules involved in transcription or whether it has a regulatory role on controlling cell fate by interacting with the genome. The impaired expression of endogenous MYOD1 mRNA in SMN-depleted cells and the temporal diffuse nuclear localization of SMN during myogenesis (**Figures 2A, S2C and 4E**) prompted us to test the hypothesis that the autoregulation of MYOD1 promotor is controlled by SMN. Indeed, chromatin immunoprecipitation-quantitative PCR analysis (ChIP-qPCR) revealed that SMN bound to the upstream region of the transcription start site (TSS) of MYOD1 in both C2C12- and iPSC-derived myogenic cells (**Figures 5A and S5A**). Furthermore, we found that SMN and MYOD1 were co-immunoprecipitated with each other in both cell types (**Figures 5B and S5B**). A physical interaction between SMN and MYOD1 was also confirmed by the overexpression of Flag-tagged MYOD1 and His-tagged SMN in HEK293 cells (**Figure S5C**). Overall, our data suggest that SMN binds to the promoter region of MYOD1 interacting with MYOD1 and regulates the expression of MYOD1 during myogenesis.

**Figure 5.**
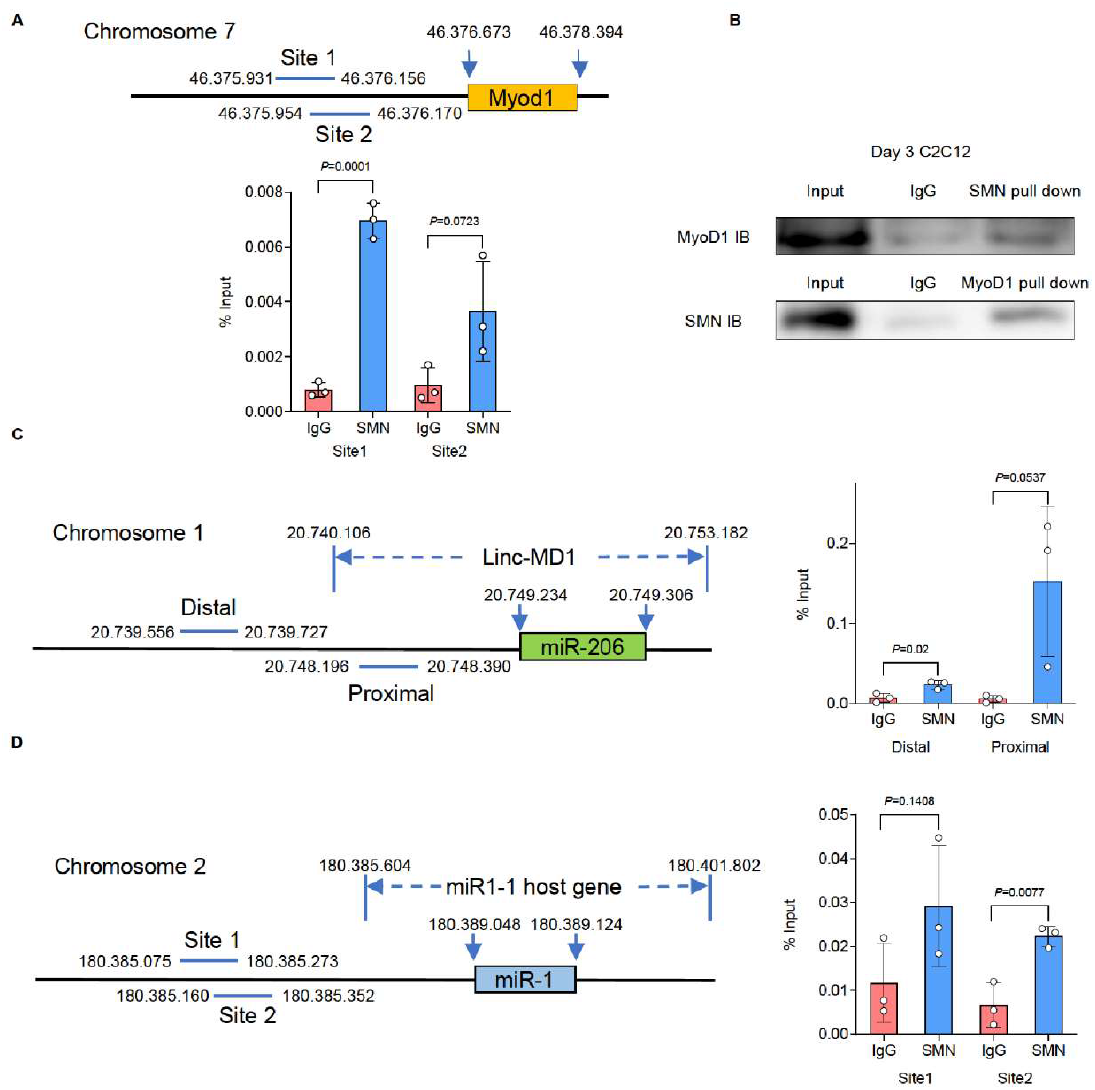
SMN binds the upstream region of MyoD1 and miRs in C2C12-derived myogenic cells. **(A)** ChIP-qPCR analysis of the SMN binding sites upstream of the MyoD1 TSS. C2C12 cells (day 3) were subjected to the analysis. **(B)** Co-immunoprecipitation assay of MyoD1 and SMN with C2C12 cells (day 3). **(C, D)** ChIP-qPCR analysis of the SMN binding sites upstream of (C) miR-206 and (D) miR-1 in C2C12 cells (day 3). Horizontal blue bars indicate the target sites. The error bars indicate means ± SD. Each dot represents a biologically independent sample.

Since the transcription of pri-miR-1 and pri-miR-206 is regulated by MYOD1 (Cesana et al., 2011; Rao et al., 2006) and because SMN and MYOD1 interact at the promotor region of MYOD1, we next investigated whether SMN also binds to the promotor region of the host genes. Linc-MD1 has two promoter regions, a distal promoter and proximal promoter, in its locus (Cesana et al., 2011). The binding of SMN in both promoter regions was confirmed by ChIP-qPCR analysis (**Figure 5C**). SMN also bound to the promotor region of miR1-1 hg (**Figure 5D**). Additionally, we confirmed that SMN binds to the promoter region of human miR-1 and miR-206 in 201B7-derived myogenic cells (**Figure S5D and S5E**). However, since SMN also binds with RNAPLII (Zhao et al., 2016), our data cannot exclude the possibility that SMN non-specifically binds the loci of other expressed genes along with RNAPLII. To exclude this possibility, we evaluated the binding of SMN and RNAPLII to the promoter region of other genes including inhibitor of DNA binding 3 (Id3), a gene highly expressed in proliferating myoblasts, and ribosomal protein L11 (Rpl11), which is ubiquitously expressed (Bouakaz et al., 2006; Wu and Lim, 2005) **(Figure S5F)**. RNAPLII bound the promoter regions of both Id3 and Rpl11, but SMN bound to neither **(Figure S5G and S5H)**. Consistently, the expression level of Rpl11 was not affected by the expression of Smn (**Figure S5I**). On the other hand, Id3 was slightly down-regulated in C2C12^siSmm^ despite SMN not binding to its promoter region **(Figure S5I)**. We attribute this observation to the disturbance of myogenic differentiation caused by the depletion of SMN (Bricceno et al., 2014). Therefore, SMN binds selectivity to certain genes during myogenesis, and SMN-bound genes are strongly down-regulated in SMN-depleted cells. Overall, SMN and MYOD1 interact with each other to bind to the promoter regions of MYOD1 and miR host genes. This mechanism may be important for regulation of the MYOD1-miR-1/-206 axis during myogenesis.

### Loss of mitochondrial integrity occurs prior to denervation in skeletal muscle of SMA model mice

Several studies have suggested mitochondrial dysfunction in SMN-deficient cells, but none have described this event in the skeletal muscle of SMA model mice. We therefore wondered whether SMN and miR- and −206 have important roles in mitochondrial homeostasis in animal models. We used a common SMA model mice, Δ7-SMA, for these experiments. Δ7-SMA mice are a transgenic strain expressing human SMN2 and the cDNA of SMNΔ7 in Smn-knockout mice (Smn^-/-^;SMN2^+/+^;SMNΔ7^+/+^). They showed an extended lifespan of 9-18 days compared to the severest SMA model mice. To understand the skeletal muscle-specific pathology, we examined Δ7-SMA mice on postnatal day 3 (P3), which is prior to the occurrence of motor neuronal deficits. The body weight of P3 Δ7-SMA mice was already decreased compared to WT, as previously reported (Ando et al., 2020) (**Figure S6A**), indicating that skeletal muscle atrophy had already begun. The expression level of sarcomeric protein actinin Alpha 2 (ACTN2) also decreased in the tibialis anterior (TA), gastrocnemius (GA) and diaphragm of Δ7-SMA mice (**Figure 6A**), which is a hallmark of skeletal muscle atrophy (Schiaffino et al., 2013). Quantification of the mitochondrial area with transmission electron microscopy (TEM) images revealed smaller mitochondria in the TA and GA of Δ7-SMA mice (**Figures 6B, 6C and S6B**). In addition, SMN protein was hardly detected in the TA or GA of Δ7-SMA mice, and mitochondrial proteins were also fewer (**Figures 6D and 6E**). We confirmed that motor neurons and neuromuscular junctions (NMJs) were maintained in P3 Δ7-SMA mice, since the areas of two neuronal markers (Tuj1 and SV2) and acetylcholine receptors (AChRs) were comparable to WT (**Figures S6C, S6D and S6E**). Therefore, the early mitochondrial defects in Δ7-SMA mice seems to be an intrinsic event of skeletal muscle preceding neuronal degeneration.

**Figure 6.**
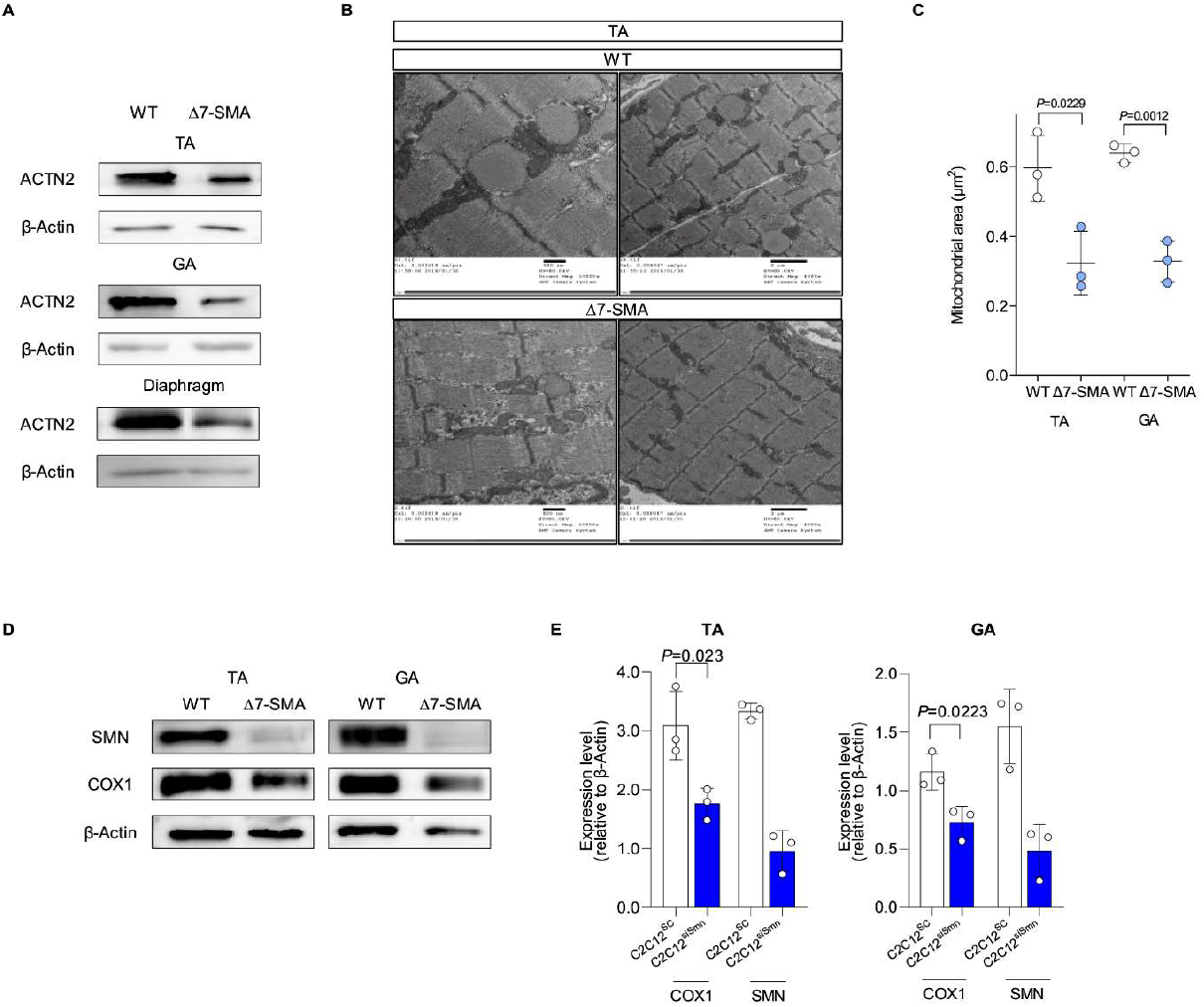
Early muscular phenotype of neonatal Δ7-SMA mice. **(A)** Immunoblotting assay of TA, GA and diaphragm samples from mice (P3). **(B)** Representative TEM images of TA muscle from WT mice and Δ7-SMA mice (P3). **(C)** Quantification of the mitochondrial area (μm^2^) in TA and GA using data from the TEM images. Three mice from each group were evaluated. The number of analyzed mitochondria is as follows: TA (WT=286, Δ7-SMA=286), GA (WT=183, Δ7-SMA=253). **(D, E)** (D) Immunoblotting assay with TA and GA samples from mice (P3) and (E) its quantification. Each dot represents independent mice. Error bars indicate means ± SD. Statistical analysis by Student’s t-test. β-Actin served as the loading control.

Severe SMA patients suffer from respiratory failure due to the paralysis of respiratory muscles (Burghes and Beattie, 2009). Considering that a loss of NMJs occurs prior to motor neuronal death in SMA model mice (Yoshida et al., 2015) and mitochondrial function is important for the maintenance of NMJs (Xiao et al., 2020), we speculated that postsynaptic mitochondria surrounding NMJs were also affected in the early life period. Postsynaptic mitochondria derived from the diaphragm of Δ7-SMA mice were significantly smaller than those from WT (**Figures S6F and S6G**). Mitochondrial proteins in the diaphragm of Δ7-SMA mice were also decreased (**Figure S6H**). Considering that the density of NMJs was maintained in the diaphragm of Δ7-SMA mice (**see Figure S6I**), these results suggested that the mitochondrial defects occur prior to the degeneration of NMJs.

### Impaired functional myotube forming ability of satellite cells from Δ7-SMA mice is recovered by miR replacement therapy

Previous data have shown that miR treatment alters the phenotype of skeletal muscle cell lines. Therefore, we next examined whether miRs could also improve the SMN-specific phenotype in mouse skeletal muscle stem cells. We applied miRNA replacement to muscle stem/progenitor cells (MuSCs) obtained from Δ7-SMA mice. For this, we isolated the cell population that included satellite cells and differentiated it into myotubes (Hayhurst et al., 2012). To evaluate the contraction ability of the skeletal myotubes, we stimulated them with a square wave electric current and measured the skeletal muscle contraction velocity (SMCV) using a cell motion imaging system (**Figure 7A**) (Hoang et al., 2019; Lin et al., 2019). Myotubes from Δ7-MuSCs showed a significantly slower SMCV than those from WT-MuSCs. Consistent results were obtained for MuSCs from both the TA and GA (**Figures 7B and 7C**). SMCV was not affected by treatment with curare, a competitive inhibitor of AChRs, excluding the possibility of NMJ-dependent muscle contraction due to an unexpected contamination of motor neurons (**Figures S7A and S7B**). Overall, Δ7-MuSCs had an impaired ability to differentiate into functional myotubes.

**Figure 7.**
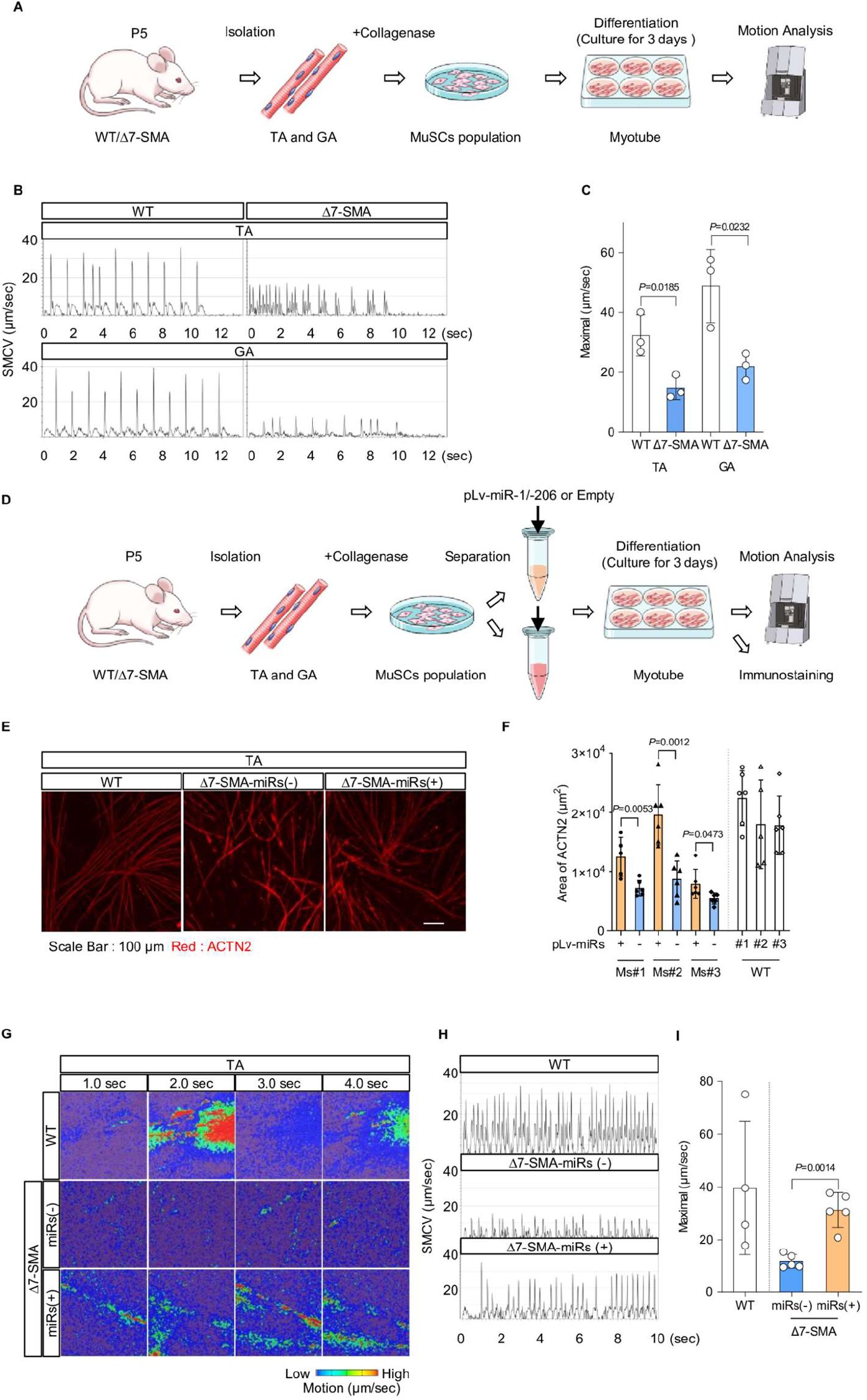
miRNA treatment improves the *ex vivo* function of MuSC-derived myotubes from Δ7-SMA mice. **(A)** Procedure for the isolation of MuSCs, myotube differentiation and motion analysis. **(B)** Motion analysis of myotubes derived from MuSCs. SMCV, skeletal muscle contraction velocity. **(C)** Maximal SMCV. Three mice from each group were evaluated. The number of analyzed ROIs is as follows: TA (WT=45, Δ7-SMA=45), GA (WT=48, Δ7-SMA=45). **(D)** The procedure for miRNA introduction into MuSCs. **(E)** Representative immunostaining images of MuSC-derived myotubes obtained from the TA. **(F)** The ACTN2-positive area (μm^2^) was calculated from (E). Three mice from each group were evaluated. Each dot represents an ROI. Six ROIs were obtained from each mouse. **(G)** Motion heatmaps of MuSC-derived myotubes obtained from the TA. **(H)** Motion analysis of myotubes derived from WT-MuSCs, Δ7-MuSCs^Empty^ and Δ7-MuSCs^miRs^. **(I)** Maximal SMCV. Four or five mice from each group were evaluated. The number of analyzed ROIs is as follows: WT=30, Δ7-SMA with miRs=31, Δ7-SMA without miRs=31. Error bars indicate means ± SD. (C, F, and I) Statistical analysis by Student’s t-test. (C and I) Each dot represents a biologically independent sample.

Finally, to evaluate the potential therapeutic ability of miR supplementation, Δ7-MuSCs were infected by lentivirus containing either an miR-1 and miR-206 double expression vector (Δ7-MuSC^miRs^) or empty vector (Δ7-MuSC^Empty^) (**Figure 7D**). Introduction of the miRs significantly improved myotube formation compared to Δ7-MuSC^Empty^ (**Figures 7E, 7F, S7C and S7D**). Notably, the SMCV of Δ7-MuSC^miRs^-derived myotubes was significantly improved and reached the level of WT-MuSCs (**Figures 7G-I and S7E-G, Video S1-S12**). Collectively, SMN plays an important role in functional muscle differentiation even in primary MuSCs, and miR replacement fully restored the differentiation potential of SMA-derived MuSCs.

## Discussion

Recently, mitochondrial metabolic dysfunction in SMA has been reported in both motor neurons and skeletal muscle (Miller et al., 2016; Ripolone et al., 2015). In the current study, we show that SMN deficiency caused mitochondrial metabolic dysfunction in C2C12 cells and myogenic cells derived from iPSCs at the early stage of differentiation. Moreover, Δ7-SMA mice showed few mitochondria and less mitochondrial protein expression at P3, which is before the occurrence of motor neuron denervation. Increasing mitochondrial ETC protein expression in C2C12^siSmn^ by miR-1 and miR-206 treatment improved the myotube formation. Therefore, our results indicated a strong association of muscle atrophy in SMA with mitochondrial metabolic dysfunction during myogenic differentiation, indicating a possible pathway for the SMA pathology in skeletal muscle. However, miR-1 and miR-206 also mediate the mitochondria-independent myogenic pathway (Chen et al., 2006; Chen et al., 2010), which was not analyzed in this study and deserves attention in future studies for its relationship with SMN.

Our results indicated that SMN could play a role in the regulation of transcription during myogenic differentiation. However, there are still questions about this mechanism. SMN protein binds with RNAPLII via the carboxy-terminal domain to form a DNA-nascent RNA hybrid structure (R loop) (Zhao et al., 2016). Failure to resolve the R loop disturbs transcriptional termination and accelerates the accumulation of DNA damage (Grunseich et al., 2018; Jangi et al., 2017; Zhao et al., 2016). However, there is no evidence that SMN deficiency in the RNAPLII complex causes the downregulation of transcripts directly. There are two hypotheses about the dysfunction of transcriptional regulation mediated by SMN on the promoter region. First is that SMN deficiency fails to form the R loop efficiently, leading to transcriptional downregulation (Grunseich et al., 2018). This failure could alter the epigenetic modification of target genes, because the R loop structure in the promoter region disturbs the binding of epigenetic modifiers. Second is that SMN could form the transcription preinitiation complex (PIC), which is essential for transcription. Thus, SMN deficiency could disturb the formation of the PIC in the promoter region of target genes. Another possible factor for the tissue-specific phenotypes in SMA is the interaction of SMN and RNAPLII. Our results showed the specific binding of SMN on the promotor region of certain expressed genes. The expression of SMN-bound genes appeared to couple with SMN binding to their promoter regions. Therefore, SMN-mediated transcriptional regulation could be one of the mechanisms for the tissue-specific phenotypes in SMA. Which factors and modifications determine the binding of SMN to promoter regions and how SMN regulates transcription are for future study (**Figure 8**).

**Figure 8.**
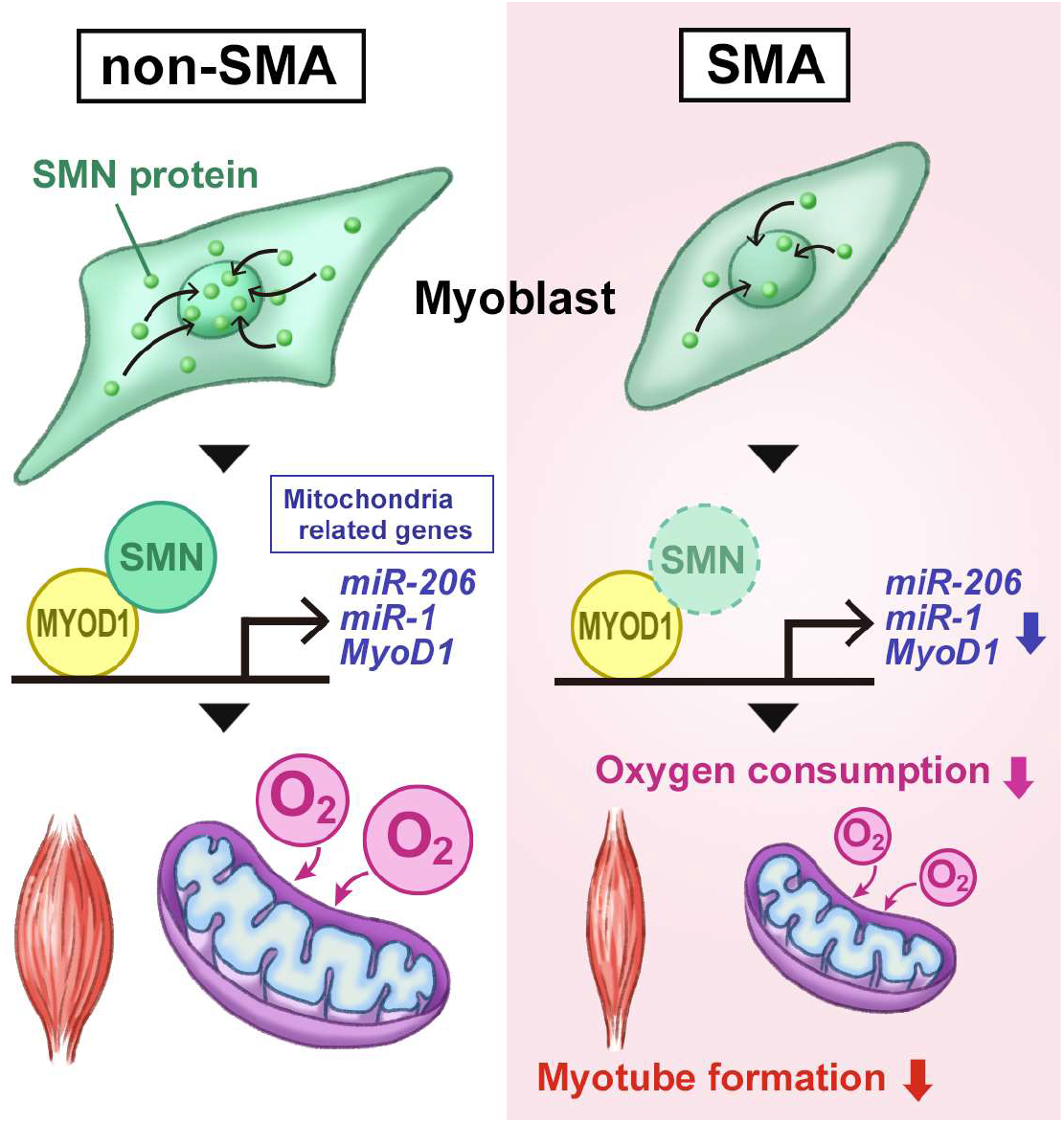
Graphical summary of the study.

We also showed that the binding of SMN at the promoter region of MYOD1, miR-1 and miR-206 regulates the MYOD1 transcriptional level. SMA motor neurons have been reported to show alternations in miR-1 and miR-206 expression (Luchetti et al., 2015; Wang et al., 2014; Wertz et al., 2016). Additionally, SMN protein binds to miR-processing proteins including fragile X mental retardation protein (FMRP), KH-type splicing regulatory protein (KSRP) and fused in sarcoma/translocated in liposarcoma (FUS/TLS) (Piazzon et al., 2008; Tadesse et al., 2008; Yamazaki et al., 2012). Therefore, it is thought that SMN deficiency disturbs miR processing, thus altering miR expression. However, there is no evidence that SMN protein has a role in the miRNA processing complex. Though we did not show whether SMN protein contributes to miR processing in myogenic differentiation, our results proved that SMN protein regulates the expression of miRs by regulating the transcription of the host genes. In our study, we found that transcriptional regulation is mediated by SMN during myogenic differentiation. Until now, there has been no report showing the SMN-mediated transcriptional regulation of motor neurons. The effects of SMN on transcription during motor neurogenesis warrant further study.

We found that the nuclear localization of SMN was transient, indicating that it binds to specific genes only. The introduction of miR-1 and miR-206 into MuSCs derived from Δ7-SMA mice improved muscle function and myotube formation. This result indicates that the postnatal introduction of miR-1 and miR-206 benefits MuSCs in SMA treatment. Supporting this finding, the miR-1 and miR-206 mediated pathway is impaired in SMN-depleted satellite cells (Qing Liu, 1997)

SMA model mice have fewer Pax7^+^ MyoD1^-^ satellite cells and a lower capacity to regenerate damaged muscle (Hayhurst, 2012; Kim, 2020), but how SMN deficiency triggers the pathology in satellite cells is unknown. Our results revealed that SMN could regulate the transcription of initial myogenic differentiation factors including MyoD1, miR-1 and miR-206. These results suggested that the disturbed initial myogenic transcriptional network by SMN deficiency could impair the maintenance of satellite cells and their capacity to differentiate. To elucidate when SMN is required and what function it has in satellite cells, we should analyze the sequential transcriptional changes in satellite cells derived from Δ7-SMA mice after differentiation and validate whether SMN nuclear localization in satellite cells corresponds to the myogenic differentiation signal.

Recently, several new drugs targeting SMN have been marketed and shown significant improvement in the prognosis of type I SMA (Mendell et al., 2017; Passini et al., 2010). However, even with early intervention with these drugs, significant functional impairment remains in the majority of cases, making it difficult to catch up with normal development (De Vivo et al., 2019; Finkel et al., 2016; Finkel et al., 2017; Mendell et al., 2017). Therefore, although therapies that increase full-length SMN expression have a significant impact on the disease course of SMA and the quality of life of patients, further functional improvement is needed to reduce the burden of the disease. One way to improve motor function in SMA patients is to develop therapies that prevent the loss of skeletal muscle in SMA patients and combine them with the restoration of SMN in motor neurons (Long et al., 2019). The regulatory function of SMN for miRs that we have discovered may contribute to the development of such a therapy. Therefore, we plan to investigate whether miR administration improves prognosis in Δ7-SMA mice.

In conclusion, the downregulation of miR-1 and miR-206 caused mitochondrial dysfunction in the skeletal muscle of SMA models. Our results indicated that SMN is the modulator for transcription during a specific phase of differentiation by binding to genome loci. Our results further suggest miR-1 and miR-206 are candidate therapeutic targets in SMA.

## Acknowledgments

We thank Ms. Harumi Watanabe for providing administrative assistance, Dr. Peter Karagiannis for proofreading the paper, Dr. Misaki Ouchida for graphical assistance, and Drs. Shiori Ando and Hideaki Hara for technical support. This work was supported by grants from the Japan Society for the Promotion of Science KAKENHI Grant Numbers 16H0552 [M.K.S] and 20H03642 [M.K.S], the Core Center for iPS Cell Research of Research Center Network for Realization of Regenerative Medicine from Japan Agency for Medical Research and Development [T.N. and M.K.S.], and iPS Cell Research Fund [M.K.S.].

## Author Contributions

A.I, M.Y. and M.K.S. conceived and designed the study. A.I. performed almost all experiments. Y.K. performed and helped with the ChIP PCR analysis. C-Y.L. performed the TEM analysis. A.N. and T. N. analyzed the data. A.I. and M.K.S. wrote the manuscript. All authors read, edited and approved the manuscript.

## METHODS

### Key Resources Table

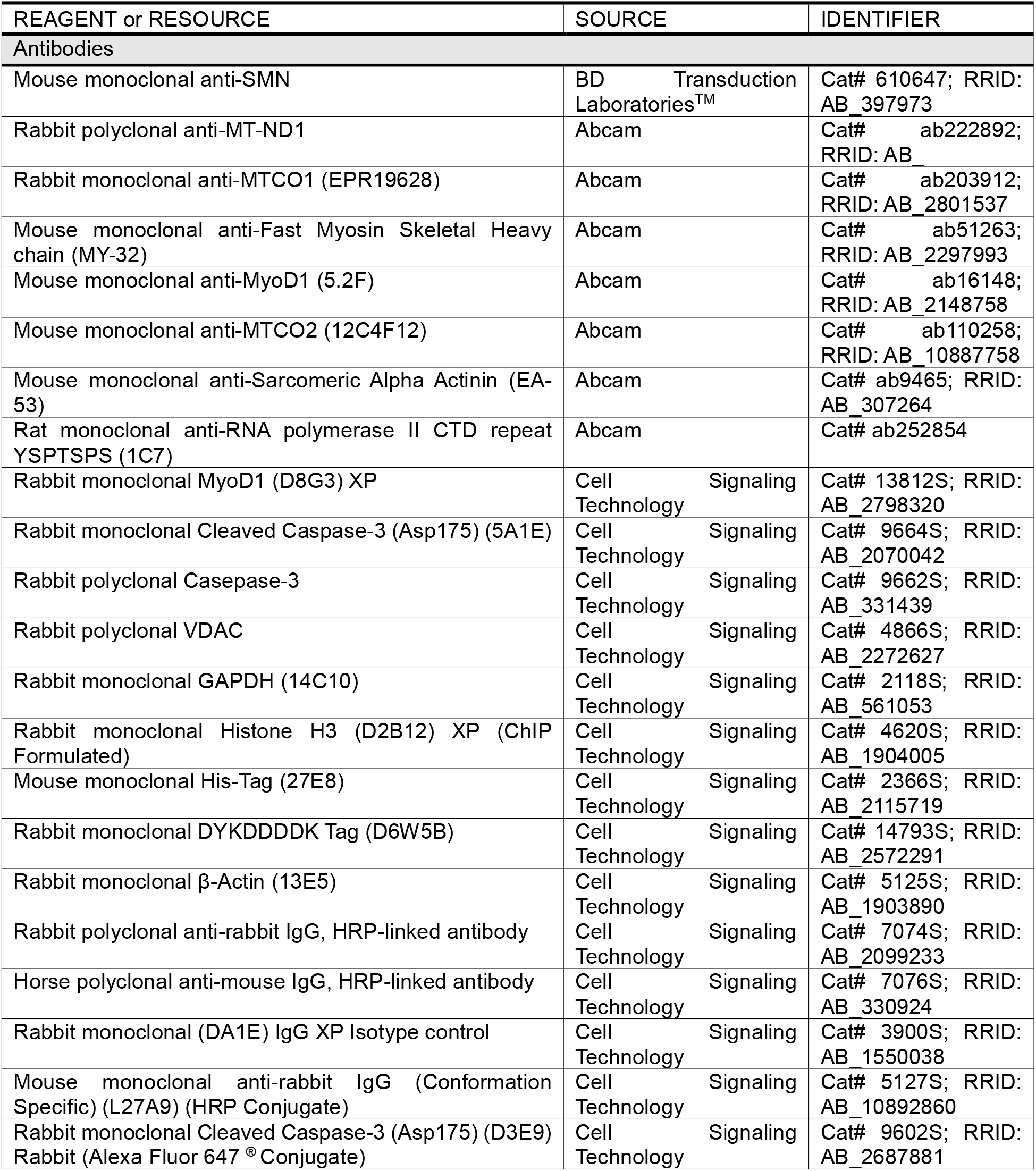
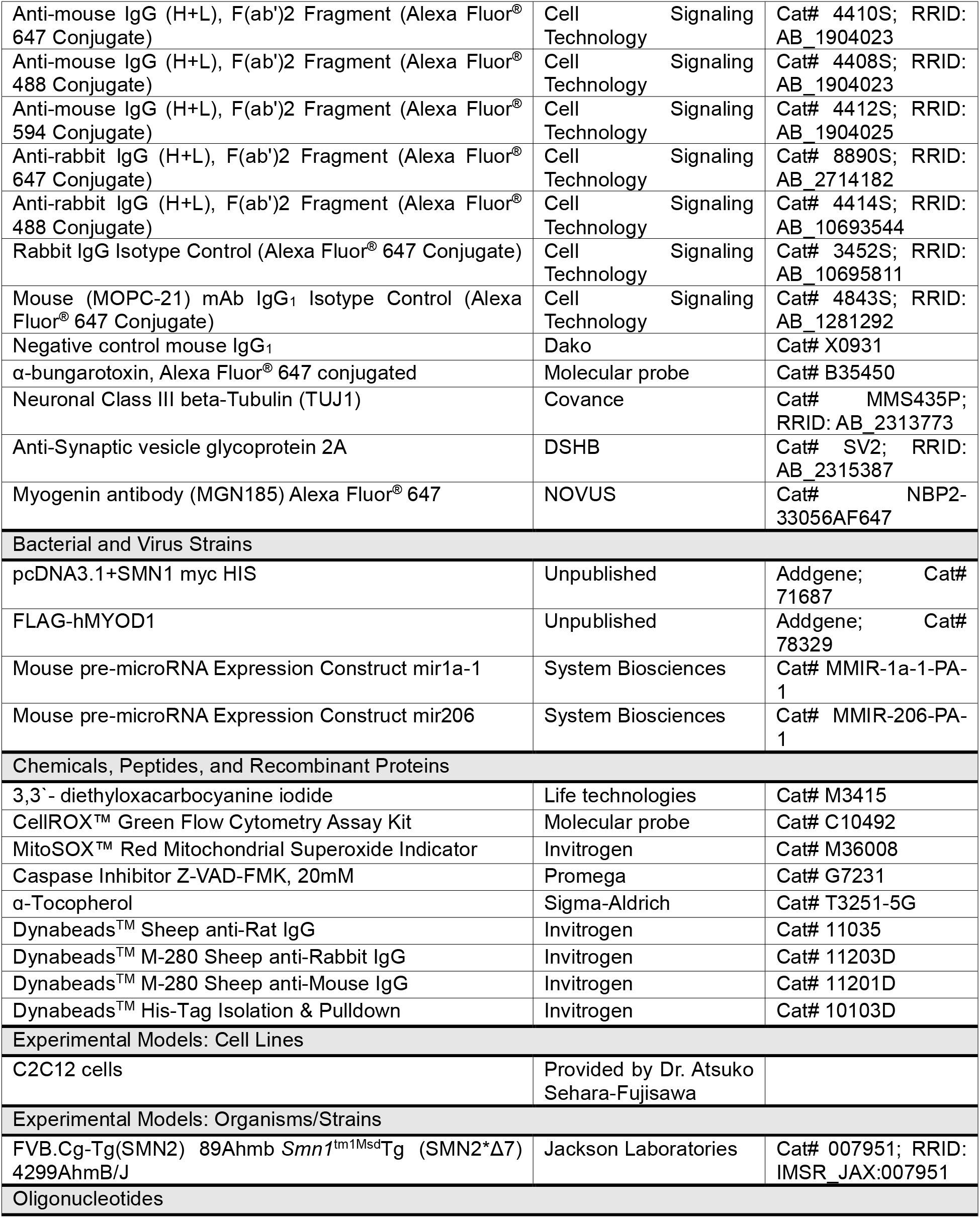
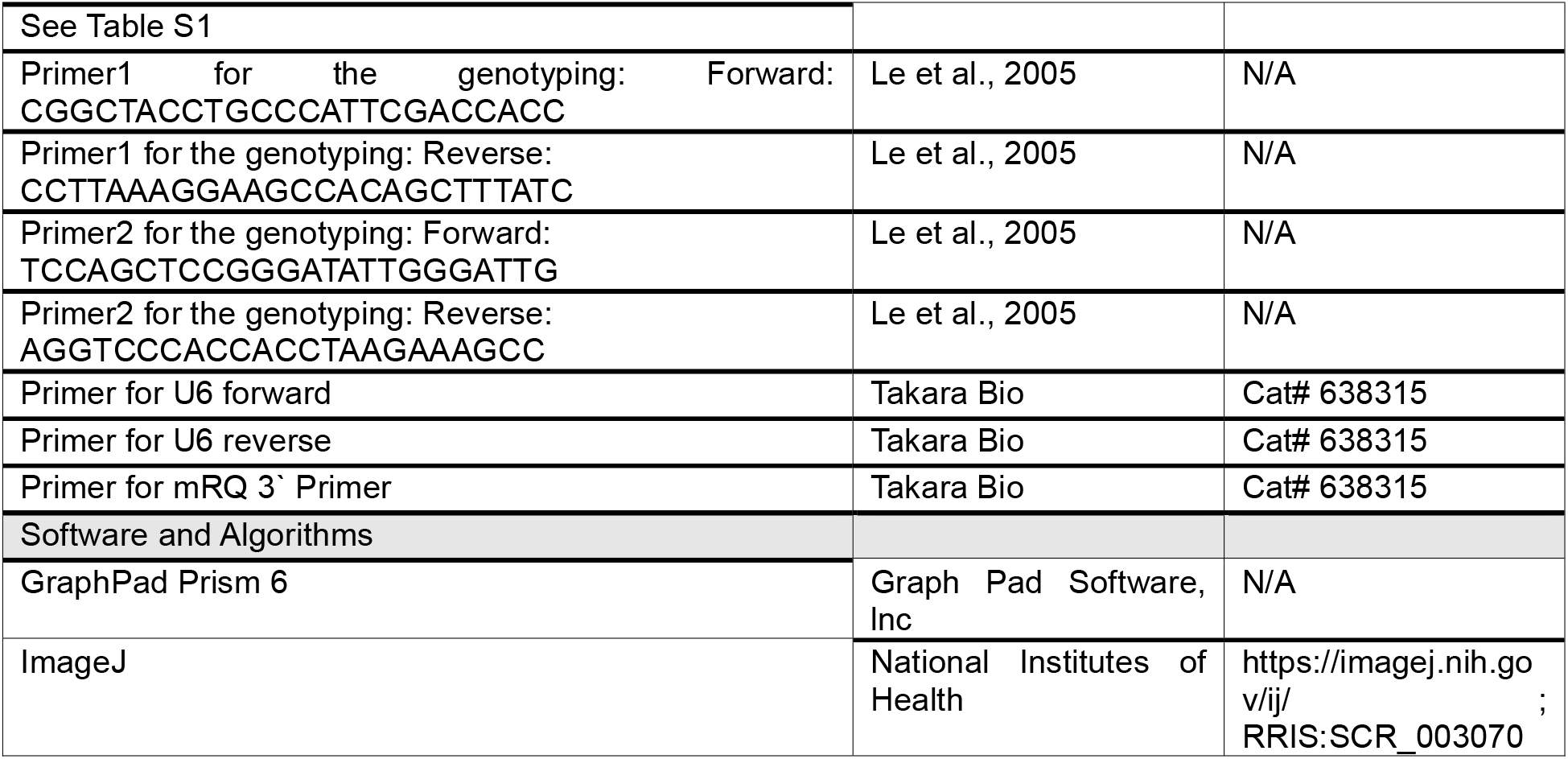

### Resource Availability

#### Lead Contact

Further information and requests for resources and reagents should be directed to and will be fulfilled by the Lead Contact, Megumu K. Saito.

#### Materials Availability

iPSCs and other research reagents generated by the authors will be distributed upon request to other researchers.

### Experimental Model and Subject Details

#### Ethics statement

The iPSC study was approved by the Ethics Committees of Kyoto University (R0091/G0259). Written informed consent was obtained from the patients or their guardians in accordance with the Declaration of Helsinki. The study plan for recombinant DNA research was approved by the recombinant DNA experiments safety committee of Kyoto University. Animal studies were approved by the institutional review board. All methods were performed in accordance with the relevant guidelines and regulations.

#### Human iPSC lines (Figure 1A)

A human iPSC line, 201B7, was kindly provided by Dr. Shinya Yamanaka (Kyoto University, Kyoto, Japan). Isogenic iPSC lines with doxycycline-inducible MYOD1 construct (B7-M and B7-M^SMNKD^) were established in a previous study (Lin et al., 2019). The iPSC line established from a type I SMA patient was also established in a previous study (Yoshida et al., 2015). The doxycycline-inducible MYOD1 overexpression vector (Lin et al., 2019) was introduced into SMA patient iPSCs by FuGENE HD (Promega) and iPSCs that stably expressed the vector (SMA-M) were selected with G418 (Wako) (50 μg/mL). A constitutive SMN1 expression vector was introduced into SMA-M to establish the SMA-M^OE^ clone, and SMA-M^OE^ was selected with Puromycin (InvivoGen) (1.0 μg/mL).

#### Conversion of hiPSC clones into myogenic cells

The iPSC lines with doxycycline-inducible MYOD1 expression vector (B7-M, B7-M^SMNKD^, SMA-M and SMA-M^OE^) were converted into myogenic cells as previously described (Tanaka, 2013). In brief, 4.0×10^5^ iPSCs were seeded onto Matrigel (CORNING)-coated 24-well plates in Primate ES Cell Medium (ReproCELL). At day 0, the medium was exchanged with primate ES Cell Medium containing doxycycline (TaKaRa) (1.0 μg/mL). From day 1, the medium was exchanged everyday with Minimum Essential Medium Alpha (Gibco) containing 10% knockout Serum Replacement (Gibco) and doxycycline (1.0 μg/mL). Cells were collected with Accumax (Nacalai tesque) on days 0, 3 and 6 after the myogenic conversion and analyzed thereafter.

#### C2C12 culture and differentiation

A mice myoblast cell line, C2C12, was maintained with Dulbecco’s minimal essential medium (DMEM) (Nacalai tesque) containing 10% fatal calf serum (FCS; Sigma-Aldrich). To differentiate the cells into myotubes, the culture medium of confluent C2C12 cells was replaced with DMEM containing 2% horse serum (HS; Sigma-Aldrich). For subsequent analysis, the cells were collected with 0.05% Trypsin-EDTA (Gibco) at days 3 and 6.

#### siRNA and microRNA mimic transfection

siSmn (Sigma, SASI_Mm01_00155410), miRNAs mimicking miR-1 and miR-206 (Gene Design) or scramble negative control (GENETOOLS,LLC, Standard control) (final concentration, 100 nM) were transfected into C2C12 cells seeded on 24- or 96-well plates using Lipofectamine RNAiMAX (Life Technologies) after the cells reached 70% confluency according to the manufacturer’s instruction.

#### RNA isolation and quantitative PCR (qPCR)

For the reverse transcription (RT)-qPCR analysis of mRNA, total RNA was extracted from the cells with the RNeasy Mini Kit (QIAGEN), and RT was performed using the PrimeScript RT Master Mix (TaKaRa). For the RT-qPCR analysis of miR, total RNA was extracted from cells with the miRNeasy Mini Kit (QIAGEN), and complementary DNA of miR was synthesized using the Mir-X miRNA First-Strand Synthesis Kit (TaKaRa). RT-qPCR was performed with TB Green Premix Ex Taq II (Tli RNaseH Plus) (TaKaRa) on the StepOnePlus Real-time PCR System (Applied Biosystems) according to the manufacturer’s protocol. Ribosomal protein L13a or U6 was used as the internal control. Primer sequences are listed in **Table S1.**

#### Intracellular flow cytometry analysis

iPSC-derived myogenic cells on day 3 were collected and incubated with 0.2% Saponin (Nacalai tesque) and 4% paraformaldehyde (Nacalai tesque) on ice for 5 minutes for permeabilization and fixation. Then the cells were labeled with antibodies against cleaved caspase-3 (1:20, CST #9602S) or Myogenin (1:20, Novus Biologicals #NBP2-33056). The following isotype controls were used: rabbit IgG Isotype Control (Alexa Fluor 647 Conjugate) (1:20, CST #3452S) and mouse (MOPC-21) mAb IgG_1_ Isotype Control (Alexa Fluor 647 Conjugate) (1:20, CST #4843S). Antibodies were incubated with cells for 90 minutes at room temperature. The labeled cells were analyzed with BD FACSAria (BD Biosciences), and the results were analyzed and processed with FlowJo software (Tree Star lnc.)

#### Measurement of reactive oxygen species (ROS) and membrane potential (Δψm)

For the measurement of the total ROS level, the cells were incubated with medium containing CellROX Green reagent (Invitrogen) (2.5 μM) for 60 minutes at 37°C. Cells were then washed three times with phosphate-buffered saline (PBS). For mitochondrial superoxide production analysis, cells were incubated with medium containing MitoSOX Red Mitochondrial Superoxide Indicator reagent (Invitrogen) (2.5 μM) for 60 minutes at 37°C and then washed three times with PBS. To measure mitochondrial membrane potential (Δψm), the cells were incubated with medium containing the 3,3’-diethyloxacarbocyanine iodide (5.0 μM) (Life technologies) for 45 minutes at 37°C and then washed three times with PBS. The stained cells were analyzed with BD FACSAria, and the results were analyzed and processed with FlowJo software.

#### Immunocytochemistry

Cells on a multi-well glass bottom dish (D141400, MATSUNAMI) were washed three times with PBS and incubated with ice-cold methanol for 15 minutes at −30°C. Fixed cells were further washed three times with PBS and incubated with 5% bovine serum albumin (BSA) (Sigma-Aldrich) in PBS for 30 minutes at room temperature. Primary antibody reactions were performed at 4°C overnight. Secondary antibody reactions were performed at room temperature for 90 minutes. Stained cells were washed three times with PBS containing 4’,6-diamidino-2-phenylindole (DAPI) (Sigma-Aldrich). Image were taken with a FLUOVIEW FV1000. (Olympus). The following primary and secondary antibodies were used: anti-SMN (1:1000, BD Transduction Laboratories #610647), anti-Fast Myosin Skeletal Heavy chain (1:1000, Abcam #ab51263), anti-Sarcomeric Alpha Actinin (1:1000, Abcam #ab9465), anti-mouse IgG (H+L), F(ab’)2 Fragment (Alexa Fluor 488 Conjugate) (1:1000, CST #4408S), anti-rabbit IgG (H+L), F(ab’)2 Fragment (Alexa Fluor 488 Conjugate) (1:1000, CST #4412S), anti-synaptic vesicle protein 2 (1:50, DSHB SV2), anti-Tuj1 (1:1000, R&D systems MAB1195), and Alexa Fluor 647-conjugated α-Bungarotoxin (0.5 μg/mL, Molecular Probes B3545).

#### Image analysis

To show myotube formation, the ratio of nuclei labeled in the DAPI-positive area to myosin heavy chain-positive area was processed and analyzed with ImageJ (NIH). To show the signal intensity of SMN in the nuclei, the nuclear area was recognized by the DAPI-positive area, and the signal intensity of SMN in the nuclei excluding SMN foci was quantified with ImageJ.

#### Immunoblotting

Cells were collected with Accumax and centrifuged at 7000 rpm for 15 seconds at 4°C. To extract proteins, cell pellets were lysed with RIPA buffer (Wako) and incubated for 30 minutes on ice. The lysate was centrifuged at 15000 rpm for 15 minutes at 4°C. The supernatant was mixed with 2 × Laemmli Sample Buffer (Bio-Rad Laboratories) containing 5% total volume of 2-mercaptoethanol (Nacalai tesque) and boiled for 5 minutes at 95°C. Polyacrylamide gel electrophoresis was performed on SDS-polyacrylamide gels, and proteins were transferred to a nitrocellulose membrane (Merck Millipore). The membrane was then incubated with 5% BSA in Tris-buffered saline with tween 20 (Santa Cruz Biotechnology, lnc.) for blocking. The primary antibody reaction was performed at 4°C overnight. The secondary antibody incubation was performed for 90 minutes at room temperature, and then the protein was detected using ECL chemiluminescence reagents (Thermo Fisher Scientific). Antibody against β-Actin was reacted for 60 minutes at room temperature. The following primary and secondary antibodies were used: anti-SMN (1:1000, BD Transduction Laboratories #610647), anti-ND1 (1:1000, Abcam #ab222892), anti-COX1 (1:1000, Abcam #ab51263), anti-MyoD1 (1:1000, CST #13812S), anti-cleaved caspase-3 (1:1000, CST #9664S), anti-caspase-3 (1:1000, CST #9662S), anti-VDAC (1:1000, CST #4866S), anti-His-Tag (1:1000, CST #2366P), anti-GAPDH (1:1000, CST #2118S), anti-Histone H3 (1:1000, CST #4620S), anti-Flag (1:1000, CST #14793S), anti-COX2 (1:1000, Abcam #ab110258), anti-Sarcomeric Alpha Actinin (1:1000, Abcam #ab9465), anti-β-Actin (1:5000, CST #5125S), anti-mouse-HRP (1:2500, CST #7076S), anti-rabbit-HRP (1:2500, CST #7074S), anti-rabbit-conformation specific-HRP (1:2500, CST #5127S), and anti-mouse-conformation specific-HRP (1:2500, Abcam #ab131368).

#### Mice

Δ7 SMA mice (Le et al., 2005) were purchased from Jackson Laboratories (FVB.Cg-Tg (SMN2) 89Ahmb Smn1^tm1Msd^Tg (SMN2*Δ7) 4299AhmB/J; stock no. 005025). Primers for genotyping were designed as described previously (Le et al., 2005). All experiments were performed on P3 or P5 Mice. Smn^+/+^; SMN2^+/+^; SMNΔ7^+/+^ was used as WT. Mice were maintained at the animal facility according to an institutionally approved protocol.

#### Isolation and primary culture of MuSCs

Isolation and culture of MuSCs were performed as previously described (Hayhurst et al., 2012). Briefly, skeletal muscle tissue was isolated from the TA and GA of neonatal mice. The muscle tissue was shredded by sterilized scissors, incubated with 0.2% (w/v) collagenase II (Roche) in DMEM containing 20% FCS and antibiotic-antimycotic (100×) (Gibco) for 30 minutes at 37°C, and dissociated into single myofibers by pipetting several times. The dissociated tissue was then centrifuged at 7000 rpm for 15 seconds and suspended with 20% FCS DMEM. The cell suspension including MuSCs was seeded into a 6-well plate coated with Matrigel. The attached cells including MuSCs were then expanded in 20% FCS DMEM condition for a few days. After reaching confluence, the MuSCs were cultured to induce myotubes in DMEM containing 2% HS for 3 days.

#### Transmission electron microscopy (TEM)

TEM sample preparation and analysis were performed following a previous protocol (Lin et al., 2019). The TA, GA, and diaphragm were dissected from Δ7-SMA or WT mice. The tissues were cut into small pieces of about 2 mm^2^ and then fixed by 0.1 M phosphate buffer (pH 7.4) including 2% PFA (Electron Microscopy Sciences) and 2% glutaraldehyde (Electron Microscopy Sciences) overnight at 4°C. Post-fixation was carried out in 1% osmium tetroxide solution (Electron Microscopy Sciences) for 1 hour at room temperature. The samples were dehydrated in graded concentrations of ethanol (30%, 50%, 70%, 90%, 95%, and 100%) and embedded in Epon resin (Electron Microscopy Sciences). Ultrathin sections (80 nm) were cut and stained with uranyl acetate and alkaline lead citrate. The specimens were examined with a transmission electron microscope (H-7650, Hitachi).

#### Motion vector analysis

Motion quantification in an ROI was performed using the SI8000 Cell Motion Imaging System (Sony) as previously described (Lin et al., 2019). Both moving image capture and motion analysis were performed using this system. To compare muscle contraction ability between WT and Δ7-SMA MuSC-derived myotubes, moving images were acquired under a continuous square wave electric current stimulation (25 V) using NEPA21 (Nepa Gene). To show muscle contraction ability, the maximal SMCV, which indicates the maximal value of the SMCV during an electric current stimulation, was applied.

#### Production and infection of lentiviral vectors

To produce the virus particle, a lentivirus expression plasmid (MMIR-1a-1-PA-1 and MMIR-206-PA-1 purchased from System Biosciences) and plasmid mixture for packaging (ViraPower HiPerform Lentiviral Expression Systems, Invitrogen) were transfected into 293 package cells with Lipofectamine 2000 (Invitrogen) under DMEM (10% FCS) condition. After incubation for 2 days, the medium containing the virus particles was recovered and concentrated with Polyethylene Glycol 8000 (Sigma-Aldrich) at 4°C overnight. The medium was centrifuged at 2000×g for 30 minutes. The virus particle pellets were suspended with PBS. For virus infection into C2C12 cells or MuSCs, virus solution was added into the cell suspension in a 1.5 mL tube, and then the mixture was incubated for 60 minutes at 37°C. After the incubation, the mixture was seeded into culture ware.

#### Immunoprecipitation (IP)

IP was performed with Dynabeads (Thermo Fisher Scientific) according to the manufacturer’s protocol. The preincubation of Dynabeads (100 μL) (M-280 Sheep anti Mouse IgG or M280-Sheep anti Rabbit IgG) with primary antibody (5.0 μL) in 1.0% BSA/PBS was performed at 4°C overnight. The beads conjugated with primary antibodies were then washed with IP buffer (10 mM Tris-HCl (pH7.8) (Nacalai tesque), 1.0% NP-40 (Nacalai tesque) and 15 mM NaCl (Nacalai tesque) EDTA free protease inhibitors (100×) (Nacalai tesque)). Cells were collected with a cell scraper in IP buffer. A total of 1.0×10^7^ cells were sonicated on ice three times for 1 second by the XL-2000 (MISONIX). Sonicated cells were centrifuged at 15000 rpm for 10 minutes. The supernatant was collected and incubated with primary antibody conjugated with Dynabeads at 4°C overnight. After incubation, the sample tubes were set on DynaMag-2 (Thermo Fisher Scientific), and Dynabeads were washed three times with IP buffer. Dynabeads were then suspended with fresh IP buffer and incubated at 95°C for 5 minutes. The supernatant was collected on DynaMag-2 and mixed with 2 × Laemmli Sample Buffer. Immunoblotting was performed as described.

#### Chromatin immunoprecipitation (ChIP) quantitative PCR (qPCR)

The preincubation of Dynabeads (100 μL) (Thermo Fisher Scientific, M-280 Sheep anti Mouse IgG) with primary antibody (5.0 μg) was performed at 4°C overnight. For ChIP-qPCR, 1.0×10^7^ cells were collected and then cross-linked in 1.0% (w/v) formaldehyde solution for 30 minutes at room temperature. Cross-linked cells were neutralized with glycine (Wako) and then centrifuged at 3500 rpm for 2 minutes. Cell pellets were suspended with 2.0% FSC/PBS, rotated for 10 minutes at 4°C and then centrifuged at 3500 rpm for 2 minutes. In the lysis procedure, cell pellets were suspended with lysis buffer 1 (50 mM HEPES buffer (Hampton Research), 140 mM NaCl (Nacalai tesque), 1.0 mM EDTA (Nacalai tesque), 10% Glycerol (Wako), 0.5% NP-40, 0.25% TritonX-100 (Thermo Fisher Scientific)) and rotated for 10 minutes at 4°C. After centrifugation at 13000 rpm for 5 minutes, the cell pellets were suspended in lysis buffer 2 (10 mM Tris-HCl (Nacalai tesque), 200 mM NaCl, 1.0 mM EDTA, 0.5 mM EGTA (Nacalai tesque)) and rotated for 10 minutes at 4°C. The cell suspension was then centrifuged at 13000 rpm for 5 minutes, and lysis buffer 2 was then replaced with lysis buffer 3 (10 mM Tris-HCl, 100 mM NaCl, 1.0 mM EDTA, 0.5 mM EGTA, 0.1% sodium deoxycholate (Wako), 0.5% N-lauroylsarcosine (Nacalai tesque)). The fragmentation of cross-linked DNA in lysis buffer 3 was performed by SFX250 sonicator (BRANSON). Fragmented DNA was incubated with Dynabeads at 4°C overnight. After the incubation, the samples were washed four times with RIPA buffer (50 mM HEPES buffer, 500 mM LiCl (Sigma-Aldrich), 1.0 mM EDTA, 1.0% NP40 and 0.7% sodium deoxycholate) on Dynamag-2 and then eluted. To detach the antibodies from Dynabeads, Dynabeads were incubated at 65°C for 15 minutes, and the supernatant was collected on Dynamag-2. Cross-linking was reversed for 24 hours at 65°C. Reverse cross-linked samples were purified using ChIP DNA Clean & concentrator (ZYMO Research) according to the manufacturer’s protocol. Eluted DNA was used as the template for qPCR. The primer sequences used for ChIP-qPCR are listed in **Table S2**.

#### Mitochondrial oxygen consumption rate

The mitochondrial OCR was measured using an XF96 Extracellular Flux Analyzer (Seahorse Bioscience) according to the manufacturer’s instruction. C2C12 cells (1.0×10^4^ cells) were seeded into XF96 Cell Culture Microplates (Agilent Technologies) coated with Matrigel and differentiated into myotubes. A total of 1.0×10^4^ iPSC-derived myogenic cells at day 5 were re-plated onto XF96 Cell Culture Microplates coated with Matrigel and analyzed at day 6. Before the analysis, the culture medium was replaced with DMEM (Sigma-Aldrich) containing 5.0 mM glucose, 5.0 mM GlutaMAX (Gibco) and 1.0 mM sodium pyruvate (Gibco). The cells were incubated for 60 minutes at 37°C without CO2. To measure the mitochondrial function, the following compounds were added: 8-10 μM oligomycin (Sigma-Aldrich), 10 μM carbonyl cyanide-p-trifluoromethoxyphenylhydrazone (FCCP; Sigma-Aldrich), and 1.0 μM antimycin A (Sigma-Aldrich) and 1.0 μM rotenone (Sigma-Aldrich). Each mitochondrial oxygen respiration-related value was calculated using Wave software (Agilent Technologies).

