## Supplementary figures for "SMN promotes mitochondrial metabolic maturation during myogenesis by regulating the MYOD-miRNA axis"

Figure S1

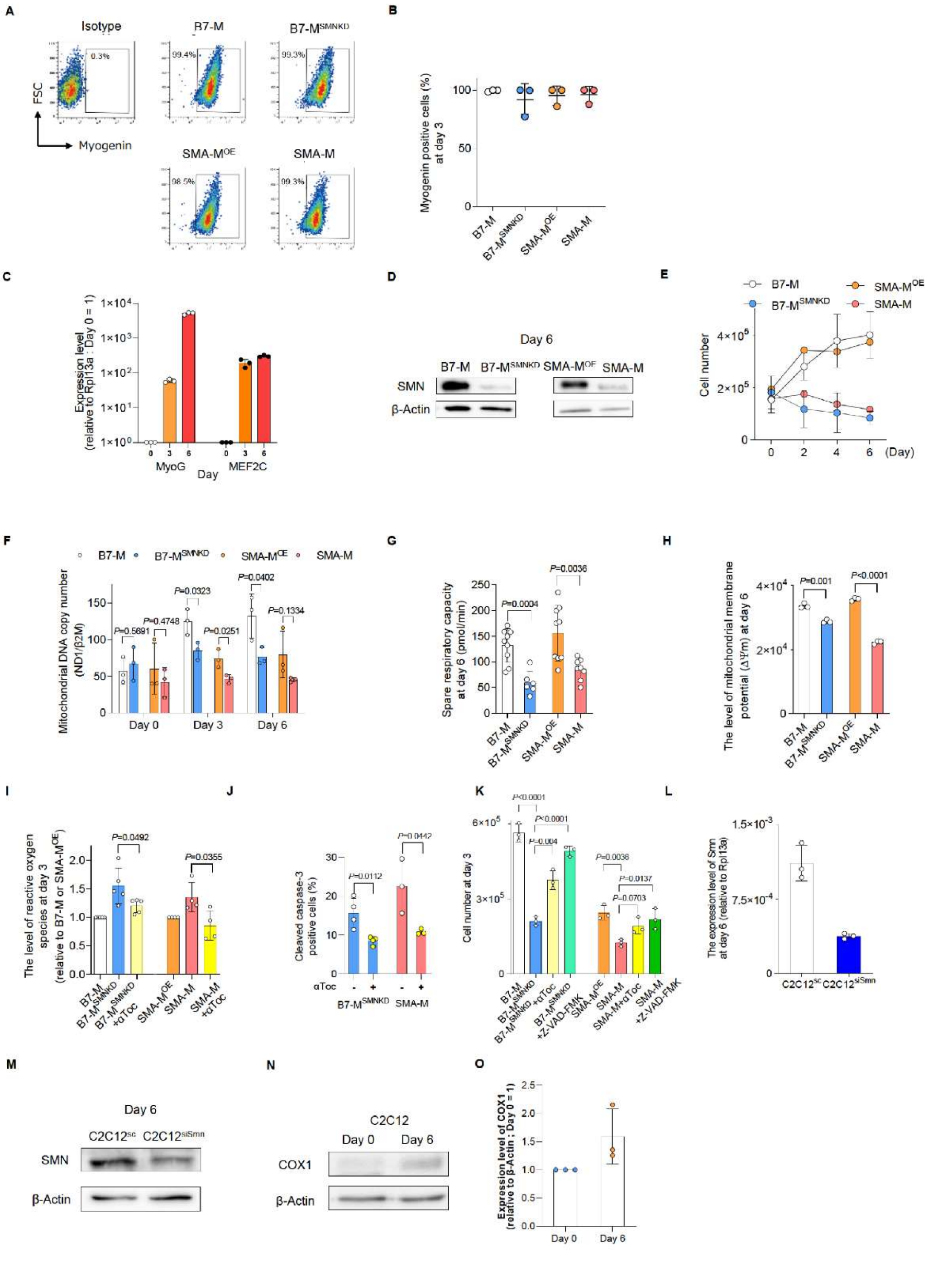

**Figure S1. Characteristics of iPSC-derived myogenic cells and C2C12 cells, related to Figure 1.**

**(A, B)** Myogenin expression in iPSC-derived myogenic cells (day 3) evaluated by intracellular flow cytometry. (A) Representative flow diagrams and (B) quantification. **(C)** qPCR analysis for the sequential expression of MyoG and MEF2C. **(D)** Immunoblotting assay with iPSC-derived myogenic cells (day 6). **(E)** Cell number during myogenic conversion. **(F)** Change in relative mitochondrial DNA copy number in iPSC-derived myogenic cells. **(G)** Spare respiratory capacity calculated using the data in Figure 1E. **(H)** Mitochondrial membrane potential ($\Delta\Psi m$) of iPSC-derived myogenic cells (day 6). **(I)** The level of reactive oxygen species in iPSC-derived myogenic cells (day 3) (relative to the MFI of B7-M or SMA-$M^{OE}$). **(J)** Cleaved caspase-3 positive apoptotic cells in SMN-downregulated clones (day 3) analyzed by intracellular flow cytometry. **(K)** Number of iPSC-derived myogenic cells (day 3). **(L, M)** Expression of Smn in $C2C12^{SC}$ and $C2C12^{siSmn}$ (day 6) at the (L) mRNA and (M) protein level. **(N, O)** (N) Expression of COX1 protein in C2C12 cells (day 6) and (O) quantification. The value for day 0 is set as 1, and relative values are shown. Error bars indicate means ± SD. (I-K) αToc, α-tocopherol (100 nM). (K) Statistical analysis by one-way ANOVA with multiple comparisons. (F-J) Statistical analysis by Student's t-test. Each dot represents a biologically independent sample.

Figure S2

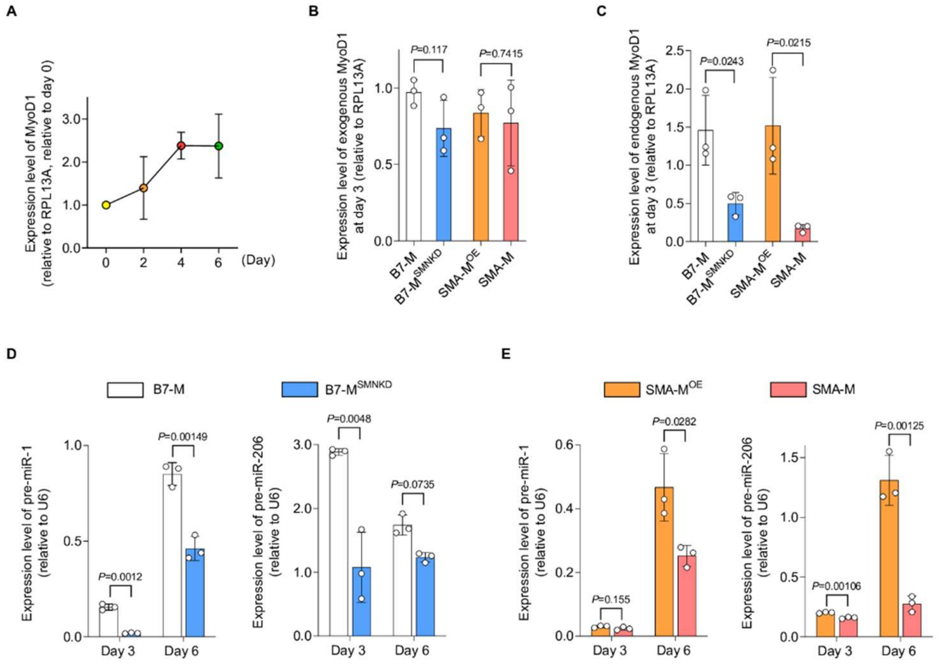


**Figure S2. Expression of MYOD1, pre-miR-1 and pre-miR-206 in iPSC-derived myogenic cells, related to Figure 2.**

**(A)** Time course of MyoD1 expression during myogenesis. Relative values to the expression level at day 0 are shown. **(B, C)** qPCR analysis for the expression of (B) exogenous MYOD1 and (C) endogenous MYOD1 in iPSC-derived myogenic cells (day 3). **(D, E)** RT-qPCR analysis for the expression of pre-miR-1 and pre-miR-206 expression in iPSC-derived myogenic cells (days 3 and 6). Error bars indicate means ± SD. (B-E) Statistical analysis by Student's t-test. Each dot represents a biologically independent sample. RPL13A or U6 was used as the internal control.

Figure S3

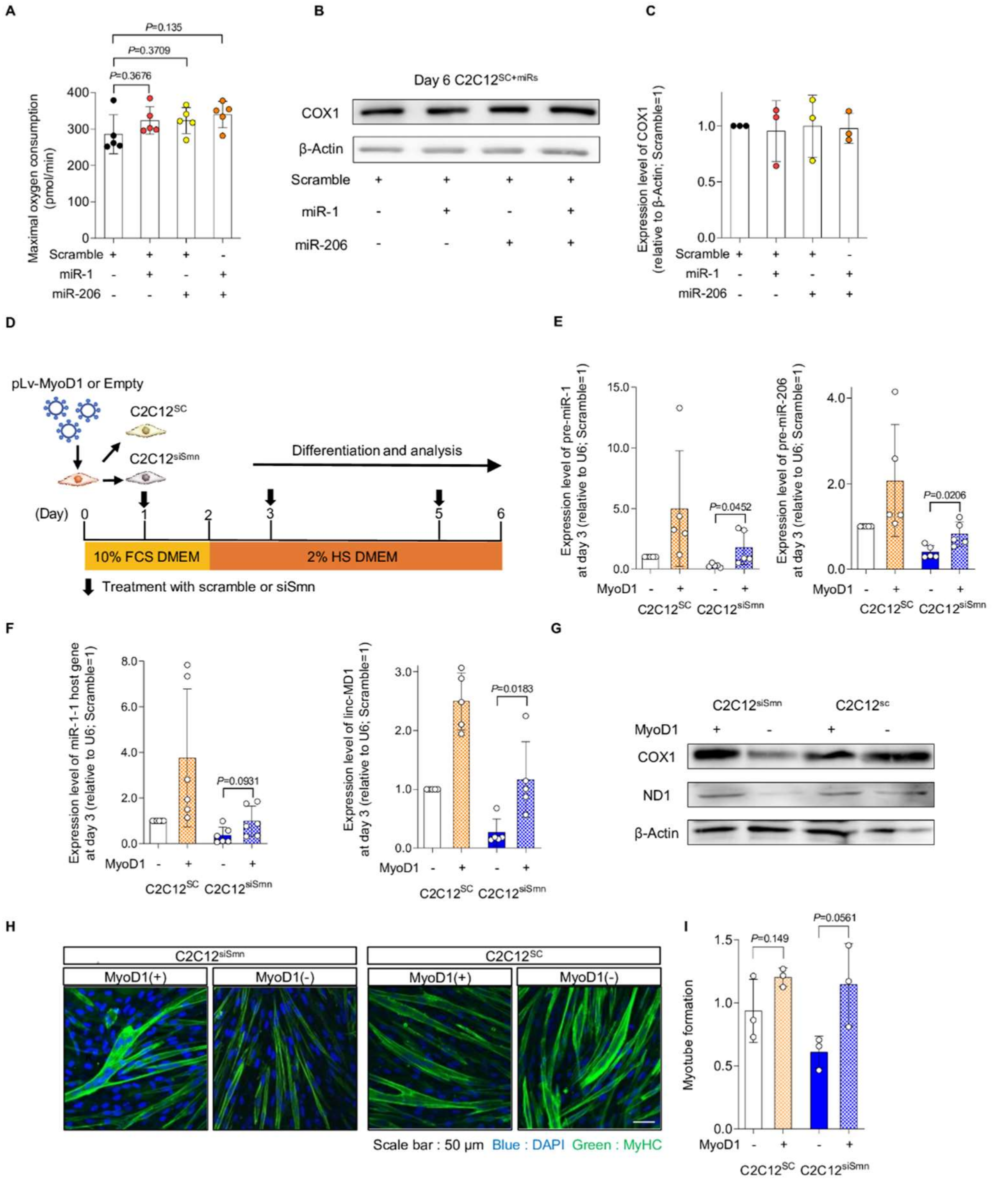


**Figure S3. Effect of miRNA treatment on C2C12$^{SC}$ and MYOD1 overexpression in C2C12 cells, related to Figure 3. (A)** Maximal oxygen consumption of C2C12$^{SC}$ cells with miRNA treatment (day 6). **(B, C)** (B) Immunoblotting assay of COX1 with C2C12$^{SC}$ cells (day 6) and (C) quantification. β-Actin served as the loading control. Relative values to untreated C2C12$^{SC}$ (day 6) are shown. **(D)** Experimental schema for the supplementation of MYOD1 in C2C12 cells. **(E, F)** qPCR analysis for the expression of pre-miRs and their host genes (day 3). U6 served as the internal control. **(G)** Immunoblotting assay with C2C12 cells (day 3). **(H)** Representative immunostaining images of C2C12 cells (day 6). **(I)** Myotube formation defined by the ratio of the

DAPI-positive area to MyHC-positive area. Error bars indicate means ± SD. (A) Statistical analysis by one-way ANOVA with multiple comparisons. (G, F, and I) Statistical analysis by Student's t-test. Each dot represents a biologically independent sample.

Figure S4

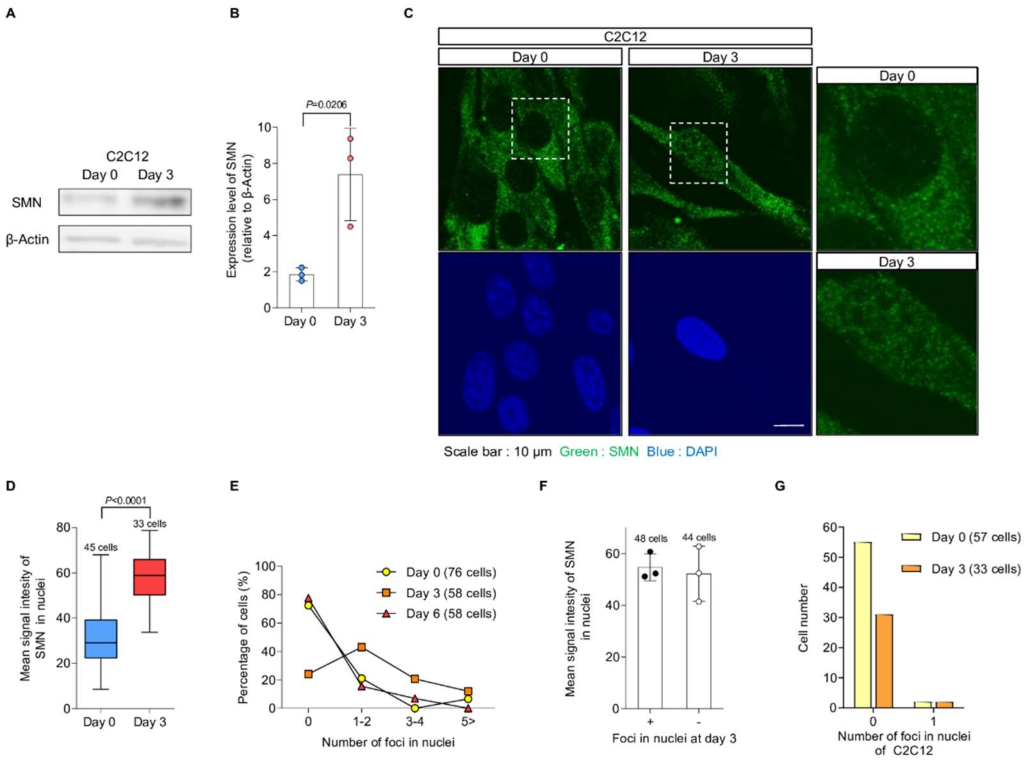


**Figure S4. Nuclear localization of SMN, relative to Figure 4.**

**(A, B)** (A) Immunoblotting assay with C2C12 cells and (B) quantification. β-Actin was used as the loading control. **(C)** Representative SMN immunostaining images of C2C12 cells. Magnified images of the dotted square regions are shown in the right panel. **(D)** Quantification of the signal intensity of nuclear SMN in C2C12 cells using the data in (C). **(E)** The distribution of SMN foci number in the nuclei of 201B7-iPSC- and B7-M-derived myogenic cells. **(F)** Quantification of the signal intensity of nuclear SMN of B7-M-derived myogenic cells with or without SMN foci. **(G)** The distribution of SMN foci number in the nuclei of C2C12 cells. Error bars indicate means ± SD. (B, D) Statistical analysis by Student's t-test. Each dot represents a biologically independent sample.

Figure S5

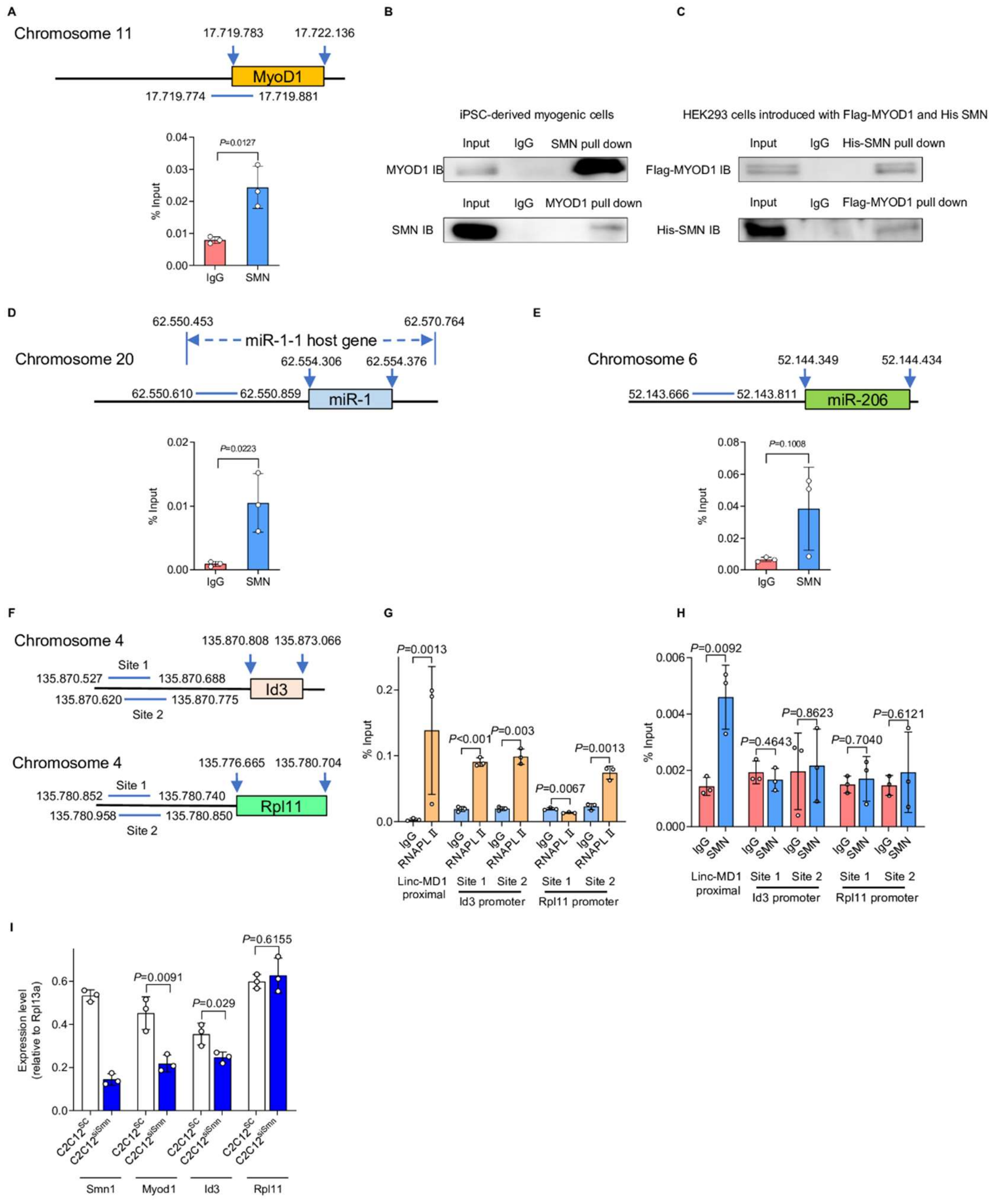


**Figure S5. SMN binds the upstream region of MyoD1 and miRs in iPSC-derived myogenic cells, related to Figure 5.**

**(A)** ChIP-qPCR analysis of the SMN binding sites upstream of the MYOD1 TSS. B7-M-derived myogenic cells (day 3) were subjected to the analysis. **(B)** Co-immunoprecipitation assay of MYOD1 and SMN with B7-M-derived myogenic cells (day 3). **(C)** Co-immunoprecipitation assay of Flag-MYOD1 and His-SMN transiently expressed in HEK293 cells. (B, C) Representative data from two independent experiments. **(D, E)** ChIP-qPCR analysis of the SMN binding sites upstream of (D) miR-1 and (E) miR-206 in B7-M-derived myogenic cells (day 3). Horizontal Blue bars indicate the target regions. **(F)** A schema of the target genomic regions of Id3 and Rpl11 for the ChIP-qPCR analysis. Blue lines indicate the target regions. **(G, H)** ChIP-qPCR analysis upstream of the linc-MD1, Id3 and Rpl11 TSS. Binding of (G) anti-RNAPLII and (H) anti-SMN in C2C12 cells (day 3) was evaluated. **(I)** RT-qPCR analysis for the expression of the indicated genes in $C2C12^{SC}$ and $C2C12^{siSmn}$ (day 3). Error bars indicate means ± SD. Each dot represents a biologically independent sample. Statistical analysis by Student's t-test.

Figure S6

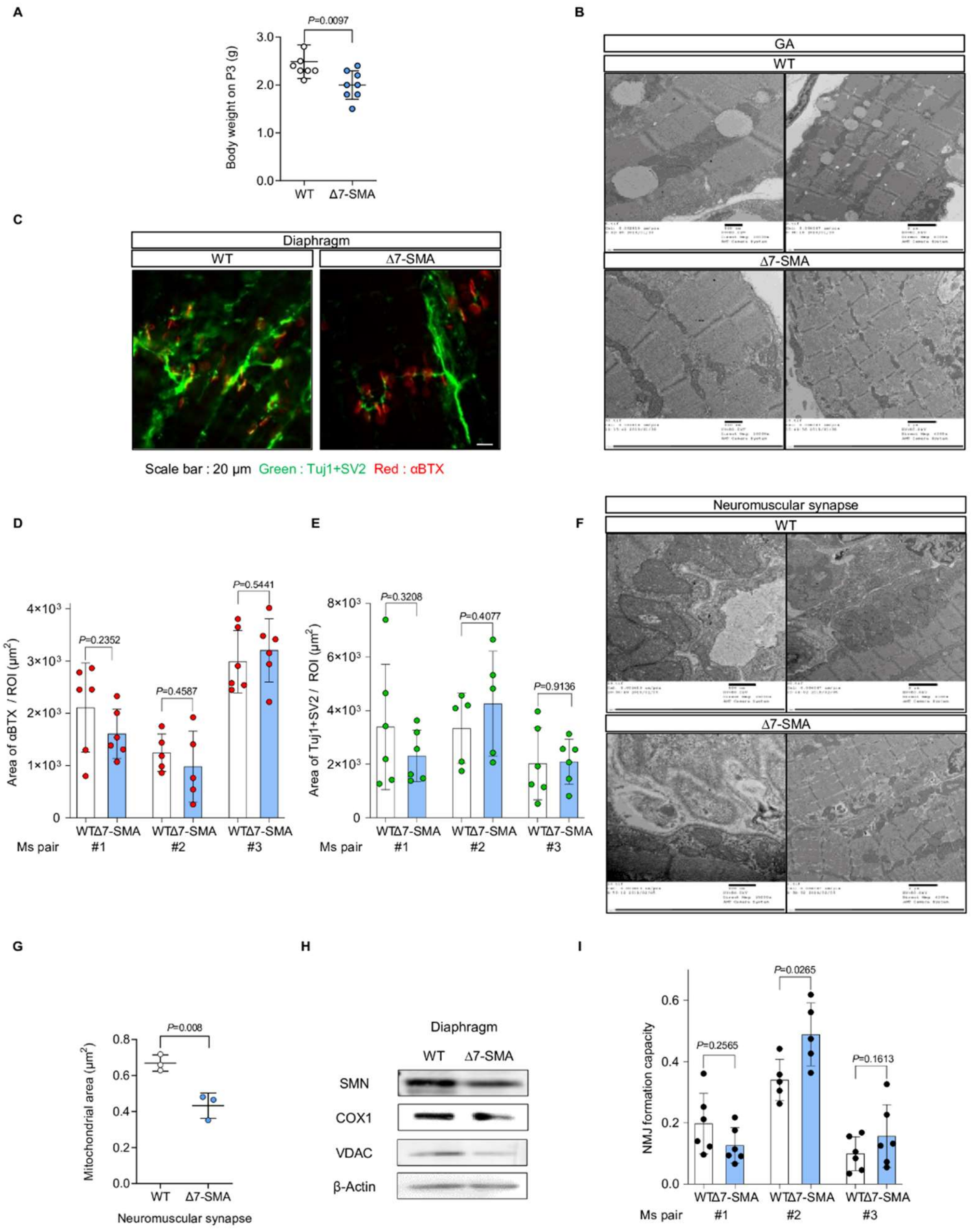


**Figure S6. Structure of the neuromuscular junction in neonatal Δ7-SMA mice, related to Figure 6.**

**(A)** Body weight of mice (P3). **(B)** Representative TEM images of the GA from WT mice and Δ7-SMA mice (P3). **(C)** Morphology of NMJs in the diaphragm of mice (P3). **(D-E)** Quantification of the αBTX and Tuj1+SV2 double-positive area using the data from the immunostaining data obtained in (C). Five or six ROIs were obtained from three mice. **(F)** Representative TEM images of the neuromuscular synapse in the diaphragm of mice (P3). **(G)** Quantification of the muscular mitochondrial area (μm$^2$) around the neuromuscular synapse using data from the TEM images. Three mice from each group were evaluated. The number of analyzed mitochondria is as follows: WT=255, ∆7-SMA=149. **(H)** Immunoblotting assay of diaphragm samples from mice (P3). **(I)** The NMJ formation capacity was defined by the ratio of the αBTX and Tuj1+SV2 double-positive area to the Tuj1+SV2-positive area. Error bars indicate means ± SD. (G) Each dot represents the number of analyzed mice. (H) Each dot represents an ROI. Statistical analysis by Student's t-test. SV2, synaptic vesicle protein 2; αBTX, α-bungarotoxin.

Figure S7

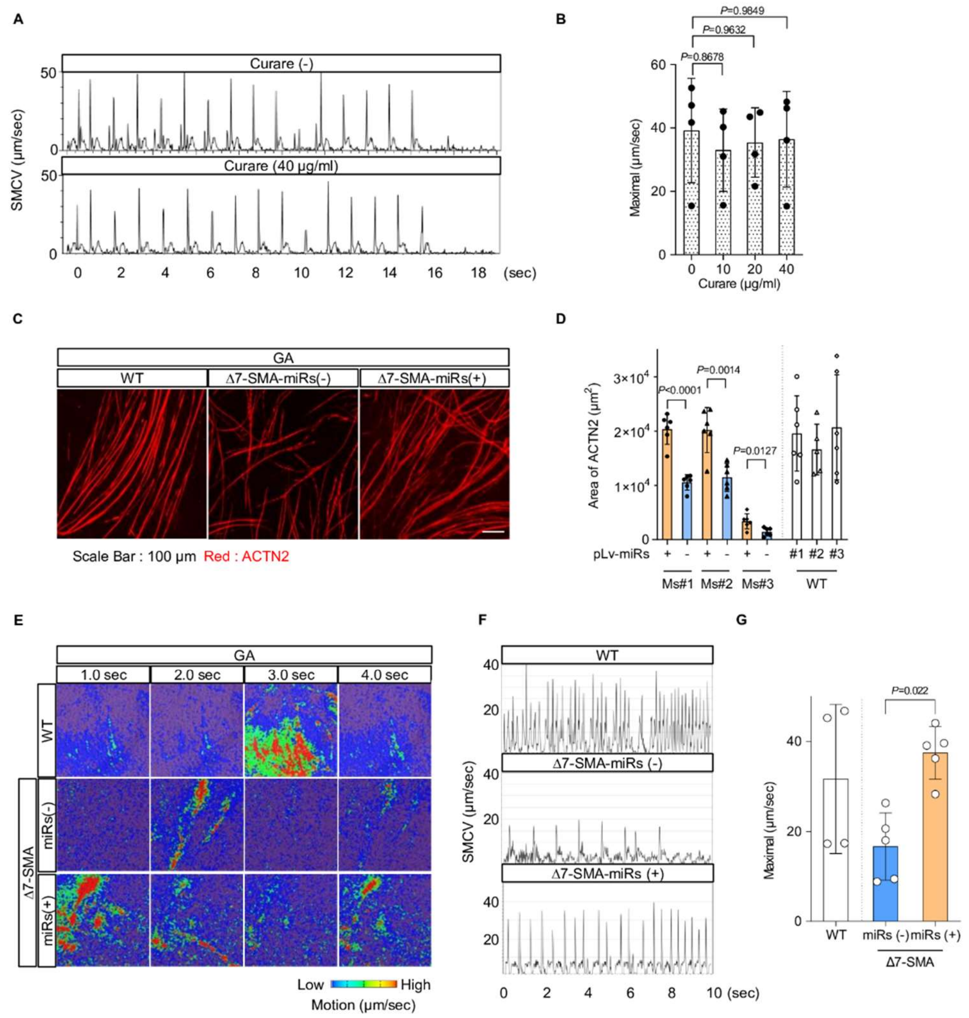


**Figure S7. Functional analysis of MuSC-derived myotubes, related to Figure 7.**

**(A)** Motion analysis of MuSC-derived myotubes obtained from the TA of WT mice with or without curare. **(B)** Maximal SMCV calculated from the data in (A). Data were from four ROIs of individual mice in each condition. **(C)** Representative immunostaining images of MuSC-derived myotubes obtained from the GA. **(D)** The ACTN2-positive area (μm$^2$) was calculated from the data in (C). Three mice from each group were evaluated. Each dot represents an ROI. Six ROIs were obtained from each mouse. **(E)** Motion heatmaps of MuSC-derived myotubes obtained from the GA. **(F)** Motion analysis of myotubes derived from WT-MuSCs, Δ7-MuSCs$^{Empty}$, and Δ7-MuSCs$^{miRs}$. **(G)** Maximal SMCV was calculated using the data in (E). Four or five mice from each group were evaluated. The number of analyzed ROIs as follows: WT=29, Δ7-SMA with miRs=38, Δ7-SMA without miRs=29. Error bars indicate means ± SD. (B) Statistical analysis by one-way ANOVA with multiple comparisons. (D, G) Statistical analysis by Student's t-test. Each dot represents a biologically independent sample.

**Supplementary video legends**

Video S1, a representative moving image capture of WT MuSC-derived myotubes, related to Figure 7G.

Video S2, a representative motion analysis movie of WT MuSC-derived myotubes, related to Figure 7G.

Video S3, a representative moving image capture of Δ7-SMA MuSC-derived myotubes, related to Figure 7G.

Video S4, a representative motion analysis movie of Δ7-SMA MuSC-derived myotubes, related to Figure 7G.

Video S5, a representative moving image capture of Δ7-SMA MuSC-derived myotubes treated with miRs, related to Figure 7G.

Video S6, a representative motion analysis movie of Δ7-SMA MuSC-derived myotubes treated with miRs, related to Figure 7G.

Video S7, a representative moving image capture of WT MuSC-derived myotubes, related to Figure S7E.

Video S8, a representative motion analysis movie of WT MuSC-derived myotubes, related to Figure S7E.

Video S9, a representative moving image capture of Δ7-SMA MuSC-derived myotubes, related to Figure S7E.

Video S10, a representative motion analysis movie of Δ7-SMA MuSC-derived myotubes, related to Figure S7E.

Video S11, a representative moving image capture of Δ7-SMA MuSC-derived myotubes treated with miRs, related to Figure S7E.

Video S12, a representative motion analysis movie of Δ7-SMA MuSC-derived myotubes treated with miRs, related to Figure S7E.
